# When attention falters: brain, breathing, and behavioral signals of lapses in interoceptive attention

**DOI:** 10.64898/2026.02.07.704566

**Authors:** Isaac N. Treves, Clare Shaffer, Alexandra Decker, Nigel Jaffe, Anna O. Tierney, Randy P. Auerbach, Christian A. Webb

## Abstract

Mindlbody practices like meditation and yoga involve paying attention to breathing sensations. During these practices, individuals report “interoceptive lapses,” moments when attention drifts away from internal bodily sensations. While lapses in attention to the external world have been widely studied, little is known about the physiological and neural mechanisms of interoceptive lapses. Interoceptive lapses may share markers with exteroceptive lapses—such as reaction time variability and default-mode network (DMN) connectivity—but may also depend on distinct brain systems and breathing physiology. Here we examined behavioral, physiological and neural signals preceding lapses in a sample of 93 adolescents enriched for GAD and depression symptoms. Participants performed a 20-minute breath counting task in the fMRI scanner with simultaneous breath recordings. Lapses were defined as moments when counting errors occurred. The sample was split into training and validation sets to test machine learning models predicting attentional lapses. The strongest predictors were timing and variability of button responses (AUCs > 0.75). Breathing variability and breathing–behavior synchronization showed smaller but generalizable predictive value (AUCs < 0.65). Whole-brain connectivity models also predicted lapses (AUC ≈ 0.65), incorporating the DMN, dorsal and ventral attention, and somatomotor networks. Further, models that included brain connectivity marginally outperformed behavior-only models. Comparisons to previous exteroceptive findings indicate some common markers (e.g., reaction time variability) and some unique markers (e.g., selective perceptual coupling with attentional networks). Although limited by the clinical sample and lack of a control task, these results highlight brain–body markers of interoceptive attention that may inform real-time monitoring during mind-body interventions.

## Introduction

Interoception, the sensing and processing of internal bodily signals, is intertwined with cognition, emotion, and behavior (1, 2). Directed attention to internal signals is a fundamental pillar in mind-body therapies like breathwork, mindfulness meditation and yoga (3, 4). Despite its central importance, researchers still have a limited objective understanding of interoceptive attention. Specifically, studies have not examined failures of interoceptive attention and how they may be predicted by behavioral, physiological, and neural changes. Objective predictors of interoceptive attention lapses could be used to track changes over the course of mind-body therapies, contribute to the development of new therapeutic paradigms (e.g., biofeedback; 5)), and even lead to passive detection of attentional failures in real-world contexts.

There has been extensive work on ***exteroceptive*** lapses of attention, or errors in attending to external stimuli. Most insights come from a limited set of laboratory tasks that involve repeatedly pressing the same button on the majority of trials, while omitting responses to infrequent stimuli (6, 7). Attentional lapses are indexed by errors in responding in these tasks (related to but not identical to the broader construct of mind-wandering, 8). Reaction time variability is a prominent indicator of attentional failures and is often higher before errors (9, 10). Respiratory patterns have also been linked to attentional performance generally, with findings indicating decreased respiratory variability in stable attentional periods (11, 12), as well as differences in behavioral performance based on respiratory phase synchronization (13, 14). For example, reaction time may be optimized when stimuli are presented during inhales and responses are initiated during exhales (14, 15). Lastly, large-scale brain networks like the dorsal attention network (involved in attention shifting, 16), frontoparietal network (involved in goal-directed attention, 17) and default mode network (DMN; involved in self-referential thought, 18) are implicated in attentional failures. DMN activation may peak preceding lapses (19), and positive connectivity between the default-mode-network and frontoparietal network (involved in sustaining attention) may precede lapses (20). Interestingly, the somatomotor network, containing well-established representations of body parts, may be linked to spontaneous task-unrelated thoughts (20, 21). Together these findings have elucidated a multisystem account of attention in task contexts, leading to the development of new paradigms for attention training (22–24). The universality of these systems (and training approaches) to the context of interoceptive attention remains uncertain.

There is some reason to believe that interoceptive attention may function similarly to exteroceptive attention. Across individuals, exteroceptive attention may be associated with interoceptive difficulties (25). Breath-focused meditation may involve downregulating the DMN, just like exteroceptive tasks (26). Extensive evidence shows interoceptive signals like respiration timing modulate exteroceptive task performance (27–29). On the other hand, there may be differences between the two processes. Researchers have found only small transfer effects from breath training practices to external tasks (30). Interoceptive attention may involve widespread deactivations of the cortex, which has not been observed during exteroceptive tasks (31). Of course, early brain **sensory** pathways for internal signals and external signals are entirely distinct (32, 33), which could result in different attentional representations even in higher-order multisensory brain areas. To this point, interrogating interoceptive attention and interoceptive attention lapses has been limited because of the difficulty of objectively measuring lapses. While one small sample exploratory study related breath-motor synchronization to self-reported task-unrelated thoughts (34), most studies have not included objective measures of attention fluctuations. For example, cardiac interoception studies typically examine how accurately participants can detect their heartbeats while sitting at rest, compared to an exteroceptive comparison condition (35, 36). Such studies typically do not assess within-task fluctuations in attention.

In the current study, we administered a breath-counting task (37) to a sample of adolescents with anxiety and depression symptoms (**Table S1**) to study respiratory interoception, and developed a machine learning approach for investigating attentional lapses. The task involves paying attention to breathing sensations and counting breaths using button presses. Specifically, participants are instructed to register each breath using one button, and press a second button when the 9^th^ breath is counted. A third button may be used to ‘reset’ the count when the participant has lost count. Measures include correct counting (‘correct cycles’), lapses of counting (‘miscount cycles’), and meta-awareness of lapses (‘reset cycles’).

An important consideration regards whether these measures specifically index interoceptive attention lapses. This task shares properties with working memory tasks (as participants need to count cycles) and metronome response tasks (as participants provide rhythmic responses to the breath)(38, 39). Importantly, breath-counting measures are not related to working memory (*r* = 0.04, *p* > 0.05) (34). This is typically interpreted as evidence that counting itself is not cognitively taxing. Second, while breath-counting shares features with metronome tasks, it differs in that the monitored signal is internally generated and must be continuously tracked rather than externally imposed. This introduces a dependency between sensory monitoring and response generation that is absent in exteroceptive paradigms. Thus, the breath-counting task involves interoceptive monitoring and reflects real-world interoceptive contexts, e.g., meditation and yoga.

Despite the clear interoceptive context of breath counting, an examination of interoceptive and exteroceptive attentional systems involved in performance has not yet been assayed. Specifically, no previous studies have identified physiological and neural mechanisms of miscounts and resets, although one previous study found that miscounts correlate with attentional lapses in exteroceptive tasks and resets correlate with questionnaires about mind-wandering (40). These patterns suggest that different response types may reflect partially distinct processes, but their underlying mechanisms—whether specific to interoceptive monitoring or reflecting more domain-general failures (like sequencing)—remain unresolved.

Rather than assuming a single underlying mechanism, the current study adopts a discriminative approach, training separate models on each response outcome, and incorporating sensitivity measures (physiological tracking of the breath) to distinguish correct counting from accurate breath monitoring. In preregistered models, we examined the preceding behavioral, physiological and neural predictors of correct, miscount and reset cycles, and assessed whether they generalize across individuals. Predictors incorporated measures used in exteroceptive paradigms such as reaction times (button press responses), phase synchronization (button press-breathing intervals), breath variability (breathing intervals), and large-scale fMRI network connectivity. To our knowledge, no previous studies of breath-counting have examined generalizable models of lapses of interoceptive attention (see **Text S1**). The first aim of our study is to identify behavioral (button), breathing, and brain predictors of attentional lapses on the task. Second, we aim to assess how robustly they generalize across people, contributing to an understanding of their future utility for new training paradigms and/or passive sensing of attentional lapses.

## Methods and Materials

### Preregistration

The present study was preregistered at https://osf.io/fetvb. A full summary of deviations may be found in **Text S2**. To test the generalizability of the models, we incorporated a held-out test sample of participants. Additionally, while used the NBS-Predict Toolbox, in light of poor training set performance we determined that the data assumptions were overly restrictive (**Text S2**). For this reason, we decided to employ a different machine learning approach involving cross-validated random forest classification, while also reporting NBS-Predict results in **Table S1**.

### Samples

Participants included 93 English-speaking adolescents (73 females, 20 males) aged 13-18 (*m*_age_ = 15.7, *SD* = 1.66), recruited from the greater Boston area as part of an ongoing randomized clinical trial (RCT) of an app-delivered mindfulness intervention (not detailed here; NCCIH R01AT011002; https://classic.clinicaltrials.gov/ct2/show/NCT04697966). Additional demographic and clinical characteristics are provided in **Table S2**. As this is an ongoing study, participants were divided into a training set (the first 61 participants enrolled, available at preregistration) and a held-out test set (32 participants enrolled over the following year). No differences in age, race, sex, current GAD, or current MDD were observed between the sets (*p* > 0.14). Only the baseline session from the trial was analyzed here.

Eligible participants had elevated levels of rumination, as defined by a score ≥ 13 on the Children’s Response Style Questionnaire (CRSQ; (40) rumination subscale. Given the higher prevalence of rumination and depressive disorders among female adolescents, our sample is predominantly female. Participants were excluded based on the following: any diagnosis of schizophrenia spectrum or other psychotic disorder, bipolar disorder, ADHD, current chronic depression (episode > 2 years), or past year substance or alcohol use disorder. Any diagnosis of neurodevelopmental disorders (e.g., autism spectrum disorder, learning disorders, etc.) that would interfere with completion of study tasks were also exclusionary. Given the fMRI scanning session, standard exclusion criteria for fMRI scanning (e.g., pregnancy, claustrophobia, cardia or neural pacemakers, surgically implanted metal devices, cochlear implants, metal objects in the body) were also in place, along with exclusion criteria for systemic medical or neurological illnesses that impact cerebral blood flow, any history of seizure disorder or head trauma with loss of consciousness ≥ 2 minutes, and current serious or unstable medical illness (e.g., cardiovascular, hepatic, renal, respiratory, endocrine, neurologic, or hematologic diseases). Participants currently taking stimulants were excluded, while participants on an SSRI or SNRI were allowed to participate, given that their dose had been stable for ≥ 2 months. Participants with active suicidal ideation were also excluded. Due to the mindfulness intervention portion of the study, participants who were currently in regular therapy, had previous exposure to mindfulness-based psychotherapy, or had extensive (≥ 300 minutes) mindfulness/meditation experience were excluded. We obtained informed consent / assent from all participants. All procedures were approved by the Mass General Brigham IRB.

### Breath counting task

During the baseline scanner session, we administered a 20-minute breath counting task (BCT; 29). The task consists of counting cycles of 9 breaths by responding to each breath using one button, and pressing a second button when the 9^th^ breath is counted. A third button may be used to ‘reset’ the count when the participant has lost count. Participants were not told when, within each breath, to press the button (at inhale, exhale, or in between), and typical presses occurred before the peak inhalation (**Fig. 1**). Full instructions are included in the supplement. Thus, the task allows for the extraction of correct cycles, incorrect cycles or miscounts, and resets. We call these responses ‘terminations’ as they ended a cycle of breathing and pressing. The standard interpretation in the breath-counting literature is that miscounts reflect attentional lapses related to vigilance, and resets represent meta-awareness of lapses (41).

**Figure 1:**
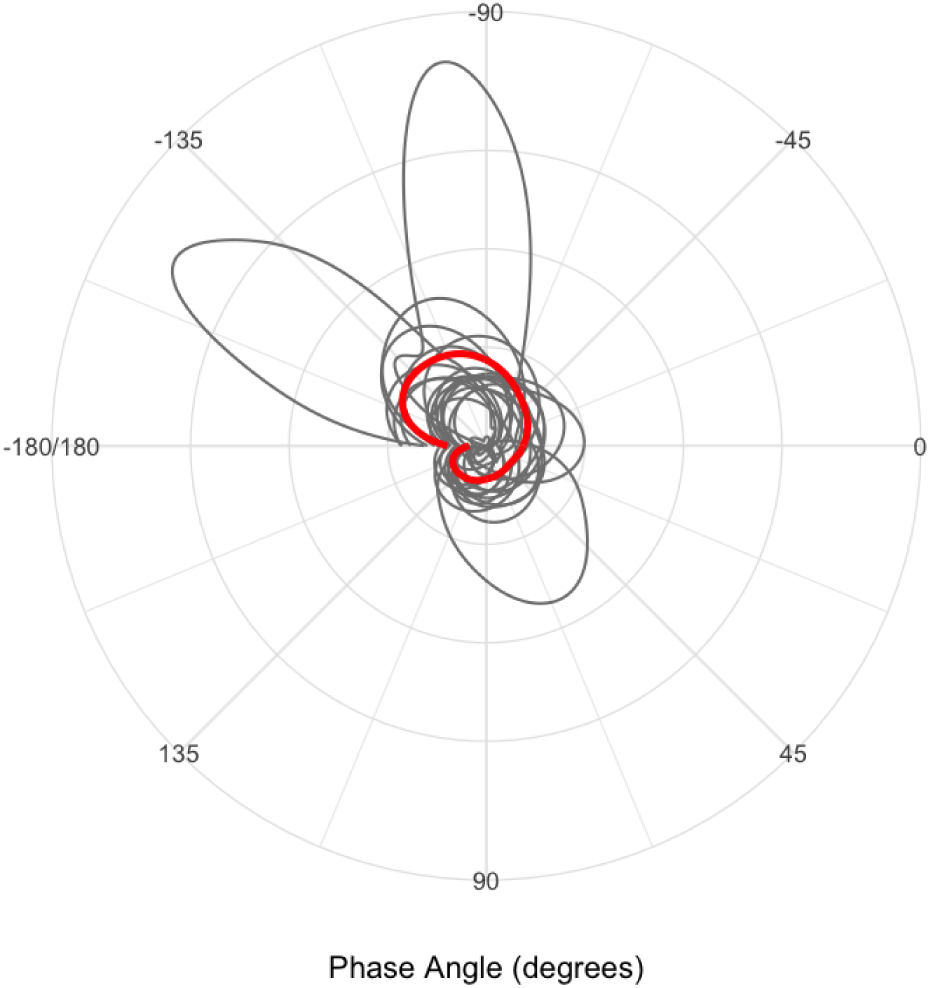
Density plots of the timing of button presses are shown for 20 random individuals (gray) as well as the group (red). Negative degrees represent button presses before the peak inhalation, where 180 represents the start of inhalation and end of the exhalation. The density reflects the proportion of breath cycles with a given timing between presses and peak inhalation. Button press timings with respect to inhalation/exhalation are included as button-breath intervals in predictive models, allowing for insight into phase synchronization and interoceptive attention.

We also collected breath belt recordings from the participants during the scan. This allowed us to a) control for breathing changes in the fMRI denoising, b) examine physiological predictors of lapses, and c) conduct interoceptive sensitivity analyses (detailed below).

#### Breath counting predictors

We extracted button press intervals, inter-breath intervals (IBIs), and breath-button intervals, all measured in seconds. Breaths were detected using peak detection software (pracma) using two peak detection algorithms ensure robustness to different breathing waveforms (for QC, see **Fig. S1**). Peaks (representing the peak inhale) were then assigned to a button using the R function *closest* which simply assigns two vectors to each other based on minimizing the distances (**Fig. S2)**. A large button-press interval reflects slow pressing, large IBI represents slow breathing, and a large absolute breath-button interval represents a long time between the peak inhalation and button press. Breath-button intervals reflect the location of responses in the inhalation-exhalation cycle and were included to test the predictive utility of measures of phase synchronization (13, 14). The three classes of predictors are not independent – e.g. for accurate performance, slow breathing necessitates slow button pressing – but there was significant variability in performance and consequentially the two were not identical (IBIs vs button-press intervals: *r =* 0.7).

### Brain imaging

#### Collection

Images were collected using a Siemens Prisma 3.0 Tesla MRI equipped with a 64-channel coil. The protocol consisted of a T1-weighted scan, three 6-minute, eyes-open, resting-state scans, a 20-minute breath-counting task scan, and then a single gradient-echo fieldmap acquisition. High-resolution structural images were acquired using a 3D T1-weighted magnetization-prepared rapid gradient echo (MPRAGE) sequence (TR = 2530 ms; TE = 3.3 ms; flip angle = 7°; FOV = 256 mm; 128 sagittal slices; slice thickness = 1.33 mm). Functional images were acquired with a multiband sequence (TR = 720 ms, TE = 30 ms, FOV = 212 mm, 66 slices, multiband accelerator factor = 6, voxel size = 2.5mm × 2.5mm × 2.5mm).

#### Preprocessing

We preprocessed the data using SPM12 (https://www.fil.ion.ucl.ac.uk/spm/software/spm12/). This consisted of realignment and unwarping using the fieldmap, slice-timing correction, co-registration of functional scans to T1w scans, normalization to the MNI template, and resampling to 2 × 2 × 2 mm voxels. Motion outliers and motion regressors were identified using ART. We then imported the unsmoothed data to CONN toolbox (42) for the final steps of preprocessing, denoising and ROI time-course extraction. In CONN, we conducted gray matter, white matter and CSF segmentation and smoothing with a kernel of 6 mm.

#### Denoising

For both resting-state and task-based (BCT) fMRI, we regressed out 5 white-matter components and 5 CSF components using anatCompCor (43). We regressed out 12 motion regressors consisting of head motion and its first-order derivatives. For the task-based fMRI analysis, we conducted additional regressions of breathing belt data (in TICS format) using RETROICOR as implemented in the TAPAS toolbox (44). Specifically, respiratory phase regressors were constructed using a Fourier expansion including sine and cosine terms up to the 4th order (eight regressors total), capturing cyclic respiration-related fluctuations. Respiratory volume per time (RVT) was additionally estimated using the Hilbert transform and included as a nuisance regressor. All physiological regressors were aligned to scan timing using scanner log files. Implementing RETROICOR and anatCompCor together provides strict control for artifacts (45–47), with the possibility of some signal loss in brainstem (48). It was not possible to control for end-tidal CO2 levels. We additionally added impulse regressors for every button press to remove motor artifacts. We did not conduct scrubbing of frames because it creates discontinuities for windowed FC analysis (49), instead we conducted despiking after regression. We bandpass filtered from 0.01 to 0.15 Hz, and conducted third-order linear detrending (50).

#### Connectivity features

We extracted timecourses from two atlases – the Schaefer cortical atlas with 300 regions and 17 networks (51), as well as the Brainnetome atlas with 280 regions including subcortical and brainstem (52). Analysis of both atlases was conducted so as to address the differential utility of the subcortex, cerebellum and brainstem. We then calculated pairwise connectivity matrices for each atlas within time-locked, HRF-delayed (6s) windows between the start and termination of breathcounting cycle. This ensures that no responses from previous cycles could influence prediction of the current termination. To control for the possibility of window length influencing performance, we conducted a sensitivity analysis with fixed windows. Results were similar (**Fig. S3 & S4**) with the exception of reset prediction, which was non-significant (**Text S3**). In addition, all main models were significant when regressing out cycle length, suggesting no overt bias on connectivity.

### Analysis

The motivation for the analysis was to develop predictive models that differentiate attentional lapses from correct cycles, consisting of training and testing models for brain, button pressing, button-breathing correspondence, and breathing independently. In addition, we identified what features drive the predictive models (timepoints during the cycles, summary features, and brain areas). Sensitivity analyses tested the performance of multimodal models, as well as whether brain features were sensitive to a different definition of interoceptive lapses.

#### Button Pressing and Breathing

This analysis deviates from standard time-locked trial-by-trial design because breath cycles and corresponding button presses are self-paced. Button press intervals, and breath-button intervals were measured at each button press, and breath IBIs were measured at each breath peak. This makes timepoint aggregation difficult as each cycle varies in breathing and button pressing. Thus, we used linear interpolation to sample the values at intervals corresponding to the TR of the MRI, 0.72 seconds, over the entire cycle. For generalizability testing, we used standard summary statistics of those values across each cycle like mean, coefficient of variation, and slope. Generalizability testing was assessed with multiple windows including the average window length of 36.89 seconds, ensuring that timepoints from previous cycles were not included. We then constructed multi-level logistic models, shown below for correct terminations, where BPI represents button-press intervals, IBI: inter-breath intervals, and BBI: breath-button intervals, and CV: coefficient of variation.

*Button-press intervals: “Correct ∼ Cycle + AverageBPI +CVBPI+SlopeBPI + Cycle|ParticipantID”*
*Breathing: “Correct ∼ Cycle + AverageIBI +CVIBI+SlopeIBI + Cycle|ParticipantID”*
*Breath-button intervals: “Correct ∼ Cycle + AverageBBI +CVBBI+SlopeBBI + Cycle|ParticipantID”*

The goal of these models was to identify which features significantly predicted binarized termination outcomes (i.e., Correct or Not, Reset or Not, Miscount or Not), while also controlling for participant and cycle number (fatigue effects). We report the significance of each coefficient estimated in the training set. To evaluate generalizability, model performance was then assessed in a held-out test set using area under the ROC curve (AUC), relative to chance. The cycle fixed effect, and random effects by participant (representing individual differences in fatigue), were removed when evaluating testing performance so as to isolate the predictive contributions of the behavioral or physiological predictors.

For testing feature importance, we deployed a time-point by time-point analysis using the average window length. To capture the nested structure of the data, we used multilevel linear models where each timepoint was predicted categorically by the termination (time was modeled categorically rather than continuously to allow for nonlinear time effects). We plotted the predicted means and 95% CIs to visually compare whether distributions for lapses vs non lapses overlapped.

#### Brain Connectivity

Random forest classification consists of training a set of decision trees (each node represents a decision point for a pairwise connectivity feature) which can be aggregated across to predict terminations. Two hyperparameters were used, leaf size and *mtry* (variables sampled), and a limited range of values was applied based on computational expense. Training consisted of assessing 5-fold cross-validated performance on the training set, combining all terminations across all individuals. This was then compared to training on 250 randomly shuffled permutations of the labels. The AUCs for the 250 permutations were sorted, and the proportion of shuffles that met or exceeded the ‘true’ AUC was compared to assess significance in the training set.

Significant models were then applied in the test set and AUC was evaluated. Additionally, head motion for each window was regressed out of the probability distribution of the classifier to assess impact on performance. Other performance metrics such as AUPRC (precision-recall), early-late fatigue effects, and individual distributions of AUCS were also calculated. For interpretation of the significant models, we assessed feature importance using out-of-bag weights and calculated the feature sums for each cortical network or subcortical area. Features sums were displayed as heatmaps on subcortical volumes, cerebellar flatmaps, and surface parcellations. We also examined the brain areas that the top 5% of feature importances belonged to by displaying positive and negative connections on a glass brain.

#### Incremental Performance of Multimodal Models

The goal was to assess whether breathing or brain connectivity may predict attentional lapses above and beyond behavioral (button pressing only) models. To test this, we conducted random forest classification using average windows (a simplification for comparability across behavior, breathing, and brain features). After removing any cycles without all three data sources (< 1%), we trained models for a) behavior only, b) breathing + behavior, c) brain + behavior, and d) brain+breathing+behavior. We then assessed their held-out performance and compared AUCs. Performance was compared descriptively where small improvements in AUC were considered ‘marginal improvements’. Other performance statistics were reported in the supplement.

#### Assessing Specificity to Interoceptive Lapses

It is possible that brain features predicting lapses are sensitive to failures of counting or working memory rather than failures of interoceptive attention, despite overall performance on the task being minimally related to working memory (37). To distinguish this possibility, we further subdivided correct cycles into *correct breathing* and *incorrect breathing* cycles. Correct breathing cycles involved correct counts and breaths (9 counts and 9 breaths), whereas incorrect breathing cycles involved correct counting to 9 *but* different numbers of breaths. This isolates counting with and without interoceptive attention, and the contrast was relatively balanced in our sample (mean correct breathing cycles = 57.0%, SD = 24.9%). Thus, we trained independent random forest models on this distinction, assessed their significance in the held-out set, and examined feature overlap with the main models (i.e., brain models that predict terminations without factoring in breathing). Overlap was calculated by conducting correlations between feature importances across models and then assessing significance through permutation testing. Finally, we examined which specific brain connections were found across both models to assess which connections may be specific to interoceptive lapses.

## Results

### Breath-counting Task Performance

Ninety-three participants performed the breath-counting task with concurrent fMRI. There was a wide range of performance on the task, providing ample variability for machine learning models. Across the sample, the average percentage of correct cycles was 56.7% (SD = 20.24%), the percentage of miscounts was 27.2% (SD=16.0%) and the percentage of resets was 16.9% (SD=15.0%). No participants were removed based on performance. Participants completed 29.7 cycles of 9 breaths on average (SD = 10.9). Breathing rates were typical for the age group(53) (M =15.11 BPM, S.D. = 3.31) and typical button presses occurred before the peak inhalation (M = -72.0 degrees, SD = 92.8°, **Fig. 1**). More descriptive statistics may be found in our previous study (41). Given the high prevalence of generalized anxiety disorder in our sample, we examined associations with task measures and model performance (**Text S4**).

### Generalizable Behavioral and Physiological Predictors of Lapses

Models were assessed for different behavioral and physiological signals preceding the terminations (correct, miscounts, resets). The strongest held-out predictive performance was observed from the model of button press intervals, or intervals between successive button presses. AUCs were 0.776, 0.744, and 0.627 respectively for correct, resets, and miscounts (**Fig. S5**; nonparameteric *ps* = .001). Other models were also significant but less predictive. Inter-breath intervals (IBIs) predicted correct and reset terminations (AUC=0.563, *p* = 0.001; AUC= 0.584, *p* = 0.001). Breathing-to-button intervals, or the intervals between the breath peak and the closest button press, predicted correct and reset terminations (AUC=0.558, *p* = 0.002; AUC= 0.618, *p* = 0.001). Miscounts were only predicted by button press intervals (AUC=0.627).

#### Discriminative features of Behavioral and Physiological Predictors

Timepoint-by-timepoint analyses of behavioral and physiological signals preceding terminations revealed substantial variability and largely overlapping confidence intervals across outcomes (**Fig. 2**). The one exception was button press intervals before a reset, which are longer for resets than all other terminations (with non-overlapping CIs). These results were largely consistent when examining different window sizes (**Fig. S6 & S7**); when not controlling for time on task (fatigue), correct cycles had shorter button-press intervals than all other terminations (**Fig. S8**). This suggests that the summary features incorporated in the predictive models were not driven by isolated timepoints. We examined the differences between the summary features by outcome (**Table 1**). The only predictive feature for breathing specifically was variability in IBIs, which was lower in correct cycles (Beta = -0.18 SD). There were large differences in button press intervals for correct cycles – less variable pressing (Beta = -0.99 SD) and faster pressing (B = -0.33 SD) was observed. As observed with the timepoint-by-timepoint analysis, resets involved longer button press intervals (B= 0.66 SD). For breathing-to-button intervals, we observed a similar relationship with variability, along with an additional effect of slope. Specifically, the absolute interval between breathing and button presses decreased over the course of a cycle for resets but increased for miscounts and correct cycles. Relationships were robust to different window lengths (**Tables S3 & S4).**

**Figure 2:**
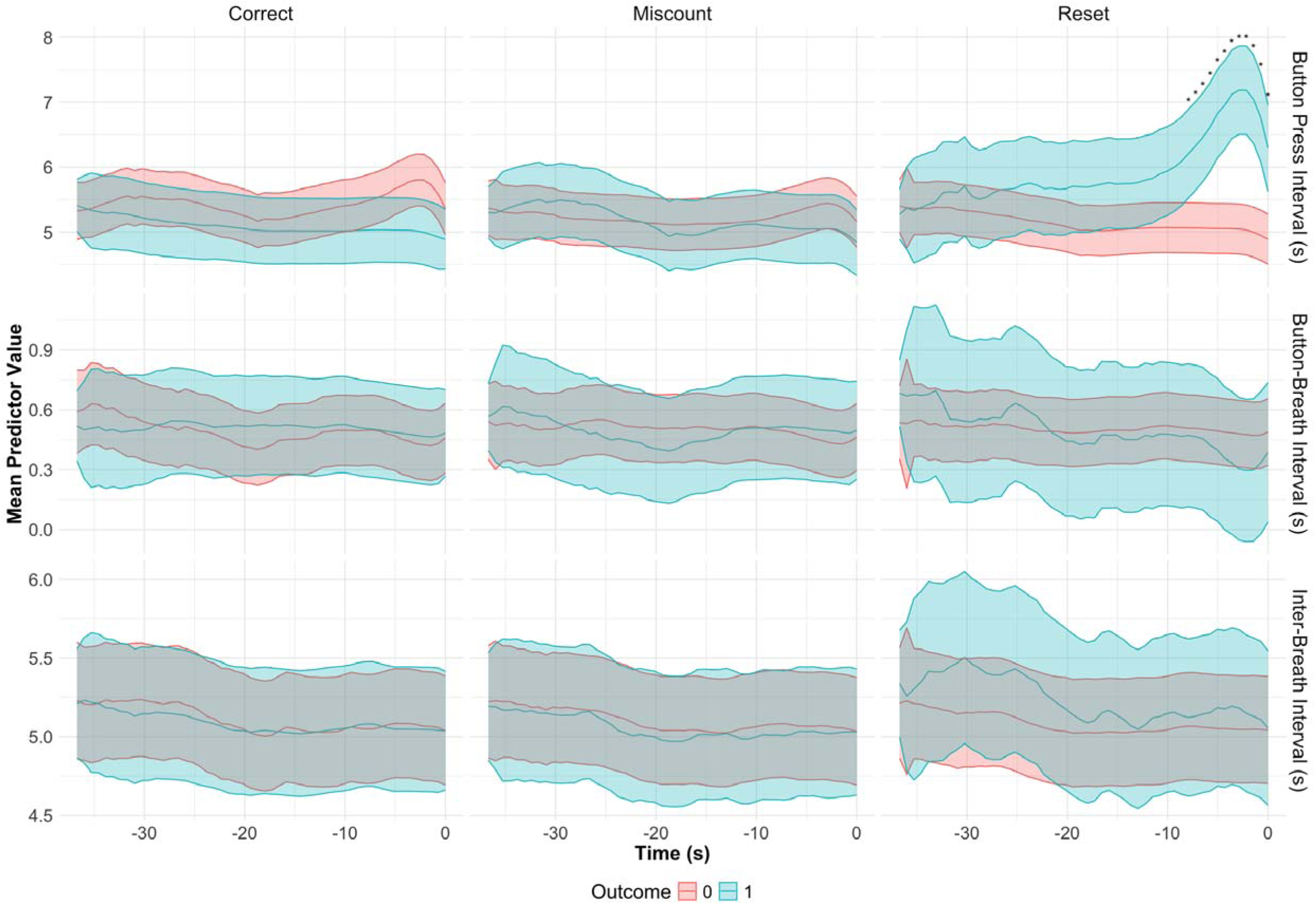
Mean predictive values of behavioral and physiological predictors are shown across individuals. Timepoints (x-axis) are shown preceding the termination of a cycle (at 0). Predictive values are shown separately for correct responses (left column), miscounts (middle column), and resets (right columns). Each row reflects a distinct type of predictor (BPI, top; BBI (middle) IBI (bottom). Blue represents the presence of the termination (e.g., ‘correct’), red is an alternative termination (e.g., not ‘correct’). Lines reflect means from multilevel models with categorical time variable to allow for nonlinearities, shading reflects 95% CI. Cycle number is included in models to account for fixed effect of fatigue. Asterisks indicate non-overlapping confidence intervals.

**Table 1:** Significant features in training set logistic models. Only features meeting strict Bonferroni threshold are displayed. Features were computed over the average time window (36.89 seconds). Betas are standardized. CV: coefficient of variation of predictor, MeanLevel: overall mean of predictor in time window, Slope_Predictor: linear slope of predictor.

| Predictor | Outcome | Feature | Effect (Beta) | P_value | 95% CI |
| --- | --- | --- | --- | --- | --- |
| Button Press Distance | Correct | CV | -0.998* | 0.0000 | [-1.162, -0.834] |
| Button Press Distance | Correct | MeanLevel | -0.333* | 0.0002 | [-0.507, -0.159] |
| Button Press Distance | Miscount | CV | 0.458* | 0.0000 | [0.332, 0.585] |
| Button Press Distance | Reset | CV | 0.442* | 0.0000 | [0.286, 0.597] |
| Button Press Distance | Reset | MeanLevel | 0.607* | 0.0000 | [0.386, 0.829] |
| IBI | Correct | CV | -0.176* | 0.0043 | [-0.297, -0.055] |
| Peak-Button Distance | Correct | CV | -0.215* | 0.0015 | [-0.348, -0.082] |
| Peak-Button Distance | Correct | Slope_Predictor | 0.202* | 0.0019 | [0.075, 0.329] |
| Peak-Button Distance | Miscount | Slope_Predictor | 0.213* | 0.0033 | [0.071, 0.356] |
| Peak-Button Distance | Reset | MeanLevel | -0.354* | 0.0041 | [-0.597, -0.112] |
| Peak-Button Distance | Reset | Slope_Predictor | -0.585* | 0.0000 | [-0.762, -0.408] |

### Generalizable Brain Predictors

Random forest models using HRF delays predicted all termination types on the task, across both brain atlases. Held-out performances, when controlling for head motion ranged from AUCs of 0.59-0.7, with *ps* < 0.001 (**Table S5).** With the exception of resets, no fatigue effects were consistently found. Controlling for the length of the connectivity window did not change results. For inspection, we have decided to focus on the prediction of correct vs miscount cycles (Whole brain AUC = 0.65, AUPRC ratio= 1.14, mean individual AUC = 0.63, 95% CI = [0.57, 0.69]), which isolates attention lapses. Important connections in the prediction of resets are shown in **Figure S9**.

#### High Importance Brain Regions and Connections

Performance was similar across whole-brain (Brainnetome) and cortical (Schaefer) models. The Brainnetome model emphasized regions of the subcortex and cerebellum (**Fig. 3, Fig. S10**). Higher connectivity predicted miscounts in visual-brainstem edges, subcortical-DAN edges, cerebellar-LIM edges, and visual-DMN edges. Higher connectivity predicted correct responses in attentional networks (ventral-dorsal attention networks (VAN-DAN),cerebellum-VAN) and within the somatomotor (SMN) network. One of the highest importance areas of the cerebellum was right lobule VIIb (**Fig. 3C),** involved in higher-order cognition including working memory. The Schaefer models highlighted attentional and somatomotor networks (**Fig. 4)**. Higher connectivity predicted miscounts in frontoparietal-visual (FPN-VIS) edges and salience (SAL)-DAN edges. Higher connectivity predicted correct responses between the DMN and SMN, FPN and SMN, and within the SMN. Notable regions implicated with a higher likelihood of a correct response included the right somatomotor areas, right insula, ventrolateral medial prefrontal cortex, the right anterior cingulate cortex, left orbitofrontal cortex, and posterior superior temporal sulcus (**Fig. 4B, Table S6)**. Notable regions implicated in miscounts included posterior parietal areas, cuneus, and middle temporal gyrus (**Fig. 4B)**.

**Figure 3:**
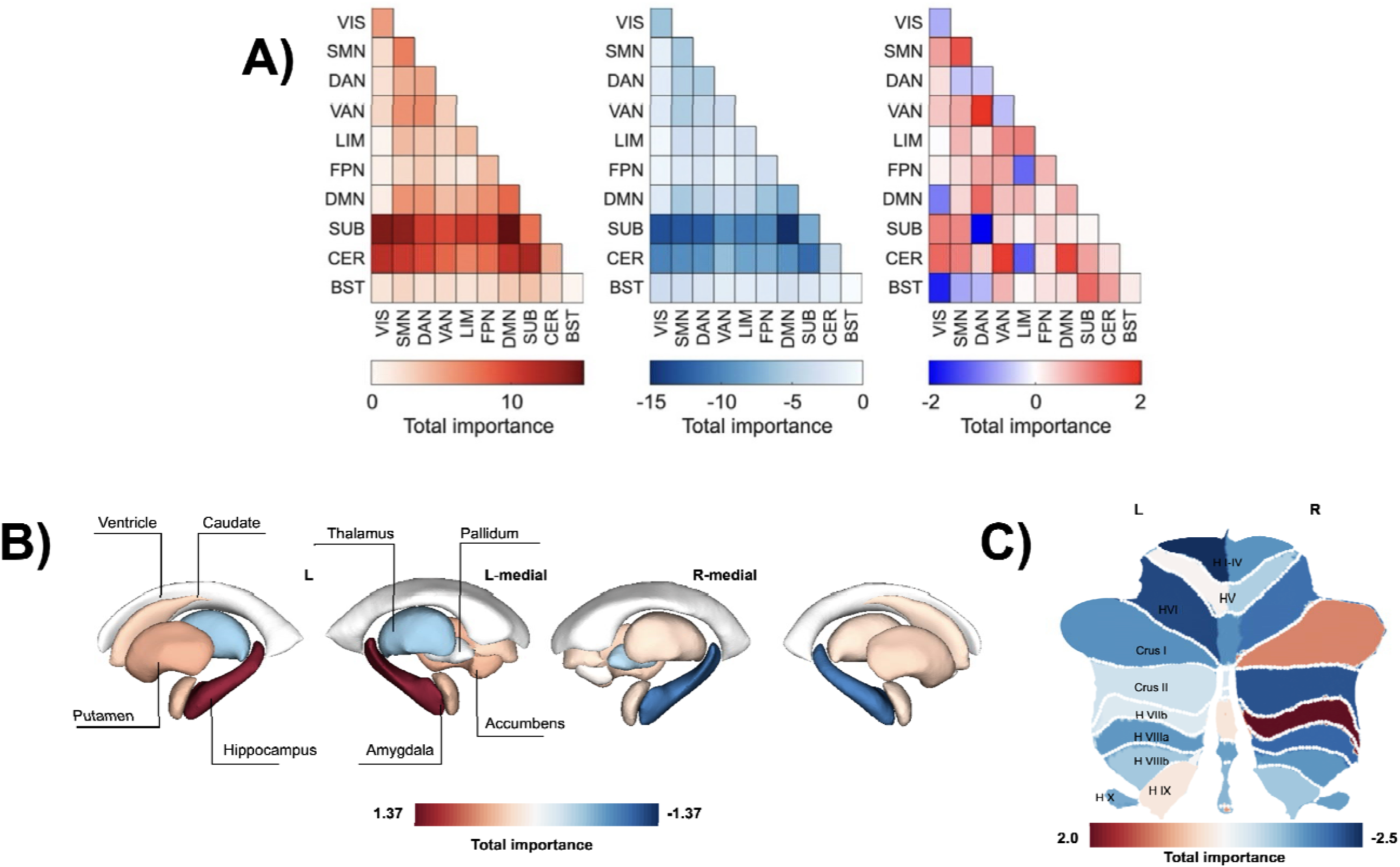
**A)** Model weights (importances) from Brainnetome models predicting correct versus miscount trials. Positive importances reflect higher likelihood of correct responses, whereas negative importances reflect higher likelihood of miscounts. Red represents positive importances, blue represents negative, and the intensity of the color reflects the total sum of importances for a combination of networks. On left, the positive importances only; middle, the negative importances; right, the total importances. B) Net importances are plotted for each subcortical parcel, which contributed substantial importance weights to the model. C) Net importances are plotted for each cerebellar parcel, which contributed substantial importance weights to the model. Blue parcels represent net negative importances, while red parcels represent net positive importance weights. Parcels in white are ∼0 net importance.

**Figure 4.**
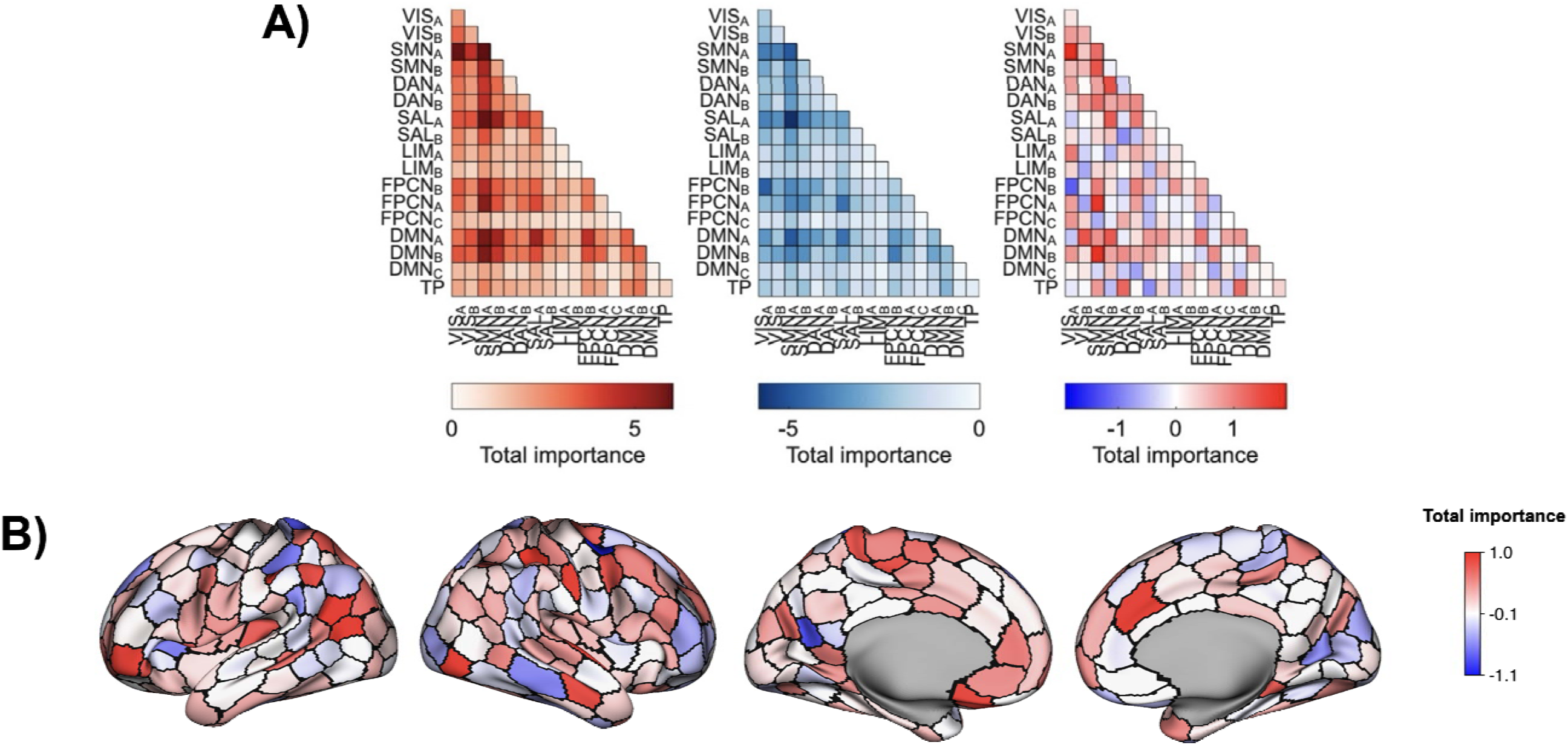
**A)** Model weights (importances) in Schaefer models predicting correct versus miscount trials. Positive importances reflect higher likelihood of correct responses, whereas negative importances reflect higher likelihood of miscounts. Red represents positive importances, blue represents negative, and the intensity of the color reflects the total sum of importances for a combination of networks. On left, the positive importances only; middle, the negative importances; right, the total importances. B) Net importances are plotted for each parcel from the Schaefer 400 parcellation. Blue nodes represent net negative importances, while red nodes represent net positive importance weights. Parcels in white are ∼0 net importance.

### Multimodal Models Perform Marginally Better

As shown in **Figure 5**, multimodal models had the highest performance in distinguishing miscounts from correct cycles (and other contrasts), but improvements over behavioral models were minor (**Table S7**). Using only mean button pressing speed, variability and change over the cycle, the models were able to achieve an AUC of 0.71. When including breathing and brain connectivity data, the models achieved an AUC of 0.75. This should be taken as evidence that some additional information is present in other modalities, but not that it is crucial to accurate prediction.

**Figure 5.**
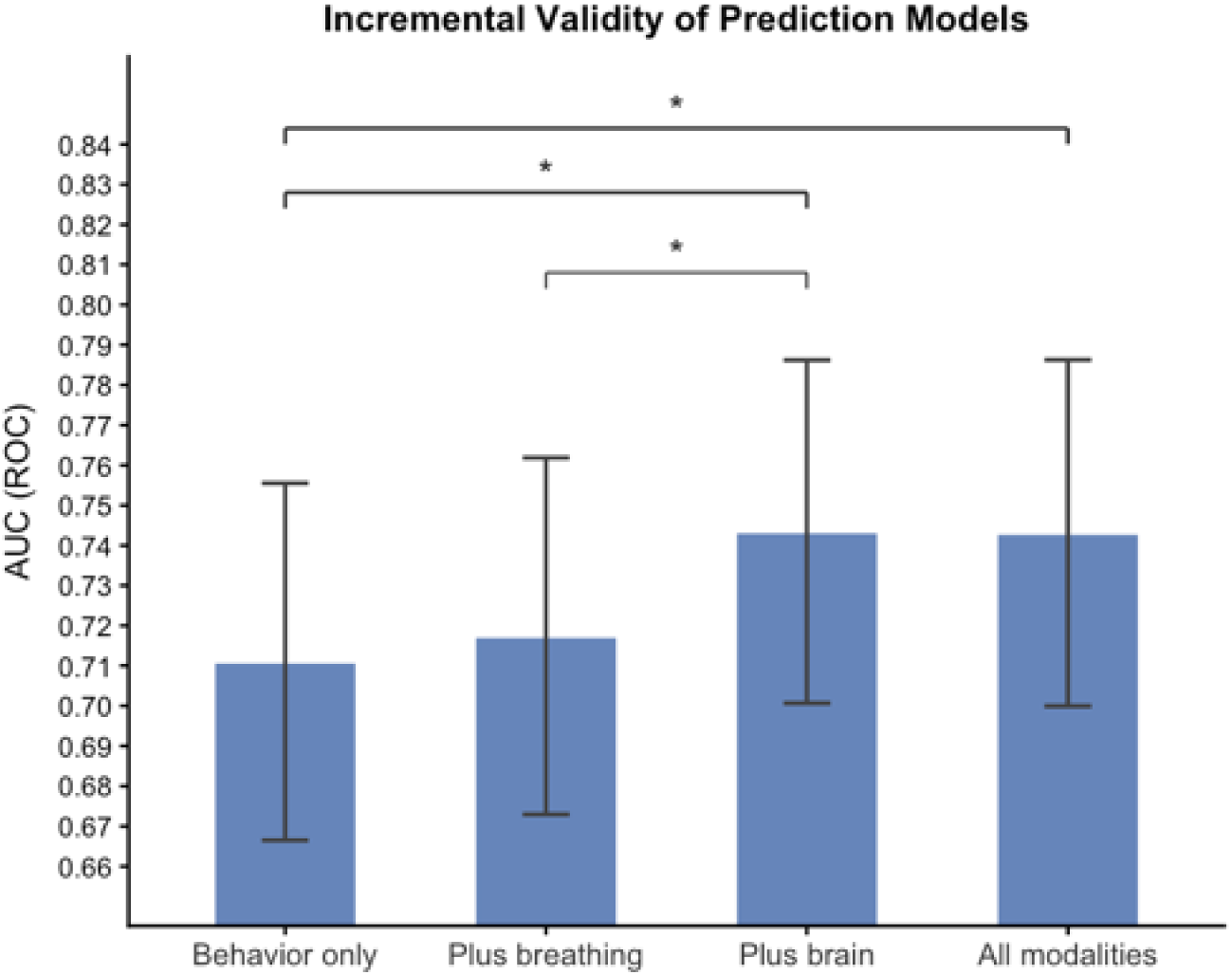
Held-out performance of multimodal models. Error bars reflect 95% CI from bootstrapped sampling. Brackets reflect pairwise one-sided DeLong tests. * *p* < 0.05. Behavior only: only button pressing features used in models; Plus Breathing: behavior + breathing features; Plus Brain: behavior + connectivity features; All Modalities: all features.

### Specificity to Interoceptive Lapses

Brain connectivity models predicting incorrect breathing (correct counting but different numbers of breath cycles) also performed significantly better than chance for both atlases (Brainnetome: AUC = 0.577, *p* < 0.01, Schaefer: AUC= 0.566, *p* < 0.01). Only the Schaefer atlas had significant overlap with the main models (*r* = 0.23, *non-parametric p* = 0.002) (**Fig. S11**). Edges that show significant overlap based on visual inspection were SMNA-SMNA, SMNA-SMNB, DAN-SMNB, FPNA-DANA and DMNA-FPNB (connections predicted correct breathing and correct counting). Edges that predicted incorrect counting and incorrect breathing included SALB-DANA and DMNC-DANB (incorrect breathing). Thus, positive connectivity in the somatomotor network may be a particularly robust indicator of interoceptive attention, whereas salience and DMN connectivity with attentional networks may be related to lapses.

## Discussion

In the current study, we leveraged multimodal objective measurement to examine behavioral, physiological and neural changes preceding lapses of interoceptive attention. Specifically, we evaluated a breath counting task (37) where ‘trials’ consisted of correct counting, miscounts (lapses), and resets (meta-awareness of lapses). Findings indicate that breathing, behavior, and brain connectivity all provide predictors of attentional state that generalize across individuals. The highest-performing model incorporated behavioral predictors (AUC > 0.75), consisting of slower and more variable button presses preceding resets and more variable button presses preceding miscounts. Reaction time variability has been robustly associated with lapses during laboratory tasks on sustained attention (9, 10). The current findings extend this literature by showing that response variability is also associated with lapses in interoceptive attention.

A novel finding from our study was that lower breathing variability (inter-breath intervals) was present during correct cycles. Additionally, breath-behavior correspondence (the intervals between button presses and the peak inhalation) also showed generalizable predictors. Notably, lower breath variability has previously been linked to periods of sustained attention during exteroceptive tasks(11, 12), and fatigue increases variability. In addition, researchers have shown that performance (response speed, accuracy, memory) is associated with the timing of responses with regards to the breath phase (13, 14, 54, 55). A previous small sample study found that the distance between pressing and breathing was associated with mind-wandering self-reports (34), but participants were explicitly told to press a button on the beginning of the inhale. In our study, which did not explicitly direct participants to respond at given times, the variability and change over time of button-breath distances was related to attentional lapses.

There is increasing evidence that there is implicit, nonconscious as well as explicit, conscious coordination between breathing and exteroceptive perceptual processes (27). Breathing is known to be linked to attentional state and arousal through the noradrenergic system and locus coeruleus. Different moments in the breath cycle are related to differences in sensory sampling and neural excitability. Thus, there are likely coordinated changes in perception across the breath cycle, as well as effects of attention on breathing (diminished variability), and our current study indicated that these changes may be detectable for predicting interoceptive lapses.

We also examined whole-brain fMRI connectivity predictors of attentional lapses. Random forest models, which generalized across participants, showed widespread networks including somatomotor, dorsal and ventral attention networks (DAN/VAN), and default mode networks. Additionally, cerebellar areas linked to cognitive control and subcortical areas involved in memory contained meaningful information(56). No evidence was found for locus coeruleus BOLD connectivity driving prediction, perhaps reflecting the poor resolution of 3T fMRI in brainstem areas. Modelling using the Schaefer cortical parcellation; however; showed that cortical areas alone were sufficient for generalizable performance. Tentatively, whole-brain and Schaefer models may have collected redundant information from cortical-subcortical and cortical-cerebellar loops involved in attentional shifting (57, 58). Networks identified in the Schaefer models were the DMN, DAN, salience (SAL), and frontoparietal (FPN) networks. While these systems are classically associated with exteroceptive attention (19–21), their interaction patterns in the present interoceptive task diverged in several key ways.

First, we observed that connectivity between attentional control networks (DAN, FPN) and the somatomotor network (SMN), as well as within-SMN connectivity, predicted correct performance, even after controlling for working memory demands. This partially aligns with prior exteroceptive findings linking attentional–sensory coupling to improved performance (20, 59, 60), often described as “perceptual coupling.” Breath perception is known to involve afferent ascending brain systems incorporating the periaqueductal gray, posterior insula (also implicated here) and somatomotor networks (33, 61). In contrast to exteroceptive paradigms where visual-attentional coupling is advantageous(62), here somatomotor–attentional coupling supported performance, whereas attentional–visual connectivity was associated with errors. Perceptual coupling appears beneficial only when aligned with the task-relevant sensorimotor processing.

Second, we did not observe the canonical pattern in which stronger DMN anticorrelation with top-down control networks predicts improved performance (often interpreted as “task-related suppression”)(20, 63, 64). Instead, distinct DMN coupling patterns dissociated correct performance from lapses: increased DMN–FPN connectivity preceded correct cycles, whereas increased DMN–DAN connectivity preceded lapses. DMN–FPN coupling may support sustained, non-evaluative monitoring of bodily signals, consistent with accounts of mindfulness-related brain plasticity(65).

While we were unable to fully exclude the possibility of condition-specific modulations in breath physiology (e.g., shallowness) confounding connectivity, our findings provide new avenues for research on interoceptive attention. Specifically, interoceptive attentional performance may depend on selective integration between DMN and control systems, alongside modality-specific perceptual coupling.

### Processes Underlying Miscounts vs Resets

A previous study used outcome correlations with other tasks to suggest that miscounts involve transient executive lapses (e.g., correlated with errors on the SART), whereas resets involve task-unrelated thought (e.g., correlated with self-report scale of mind-wandering)(66). Our study provides convergent evidence that resets are a more sustained form of interoceptive disengagement.

High-resolution tracking of behavior, breathing, and neural connectivity allows for novel, granular insight into response processes. Both miscounts and resets involve more button-press variability and more breathing variability. However, resets were specifically preceded by markedly lengthened button-press intervals in the absence of corresponding changes in breathing rate.

Interestingly, while neural connectivity can be used to predict miscounts and resets, resets were preceded by different patterns of perceptual coupling. Namely, subcortical – SMN and subcortical-VIS connectivity predicted resets, in contrast to miscounts which were predicted by less control network – SMN connectivity. This could suggest that bottom-up processes of sensory processing are present (and perhaps control-SMN connectivity differences are more transient). Interestingly, resets are not predictable when using fixed, ‘mean’ length windows. It is possible that artifactual concerns influence findings, but reset performance was significant when controlling for window length and head motion. This could suggest attention lapses emerge early before resets, and fixed windows may miss critical neural information.

While neural models showed some lack of robustness for resets, they outperformed models of miscounts when controlling for working memory confounds (Reset AUC = 0.693, Correct Breathing AUC *=* 0.577). This is consistent with behavioral and breathing models predicting resets outperformed miscounts. This finding suggests that neural models predicting sustained interoceptive vs exteroceptive engagement are more effective than neural models predicting transient executive lapses.

### Applications

There is substantial interest in developing methods for facilitating mind-body interventions like breathing meditation (67–69). The present work provides multiple indicators of attention that could be deployed to monitor or provide feedback on breathing attention. Breath variability and brain connectivity are promising as they can be collected without requiring user input. While these markers are associated with attention, the accuracies are relatively modest (AUCs < 0.7), and incremental validity above and beyond behavior is likewise modest. Models may improve with person-specific calibration, perhaps by learning from signals collected in less cognitively demanding contexts(31). Integration of non-invasive and scalable monitoring systems like OPM Magnetoencephalography could provide high temporal resolution signals for predictive models(70). Alternatively, biofeedback applications using smartphones to monitor breathing for links to attentional and affective states may be indicated (69).

A second application concerns clinical research on disorders of interoception and body awareness, including anorexia, chronic pain, panic disorders, and more. Self-report scales in these populations may indicate disturbances in interoceptive attention (e.g. higher attention in anxious populations) (2). Using just self-report, it is unclear whether the salience of interoceptive attention to their emotions and well-being is higher, or whether individuals actually spend more time attending to their bodies. Objective measures, especially non-invasive measures, may provide insight into fluctuations of attentional state in clinical populations. Particularly promising may be multimodal fusion methods (e.g. behavior-breathing correspondence). A multisystem approach could provide new insight into links between interoceptive disturbance and well-being.

### Limitations

A strength of our study was relatively long (20-minute) scans and sufficient participants for held-out tests of generalization; a weakness is that our participant sample was constrained to largely female, ruminative adolescents with a high prevalence of GAD (41.9%). Characteristics of the clinical development sample affected performance. While performance across the whole sample was below norms, an exploratory analysis of GAD individuals revealed that they performed more accurately (fewer miscounts). Moreover, model performance across modalities was better for predicting resets in GAD individuals (**Text S4**). While exploratory, this suggests sample differences in the saliency of breathing sensations and performance monitoring are associated meaningfully with response processes and mechanisms of interoceptive attention.

Second, we preregistered network-based predictive models (NBS-Predict,59), which showed poor training set performance and failed to generalize to held-out participants. We believe this was because the NBS-Predict algorithms only allowed for linear classification (e.g., SVM), and required that any predictive edges form ‘connected components’ in order to mimic brain network organization. Thus we chose to examine random forest models instead, which show promising results in other predictive modelling studies and allow more flexibility in feature selection (72, 73). Our random forest models generalized, showing significant performance (AUC = 0.65) in held-out participants (a high bar, 63). Future studies should further test the benefits and tradeoffs of ML algorithms in brain-based prediction studies.

A notable limitation of the study was that lapses on the task could have originated from failures of working memory (counting) or from failure to attend to sensations of the breath. Thus it is possible that predictors herein are related to working memory (74) and not a lack of attention to the breath per se. This concern is somewhat but not completely alleviated by the simplicity of the counting task, and observations that working memory does not robustly correlate with breath-counting performance (e.g. OSPAN; 32). In order to further address this concern, we conducted sensitivity analyses which revealed that brain connectivity can distinguish trials with similar working memory (identical counting) but differences in breathing. The connections revealed by this approach overlapped with the main models, but only in cortical areas. It is likely that our main models predict performance adequately on ‘breath counting’, but may be more limited when addressing other respiratory paradigms, for example, tasks involving detecting breathing rates or imposed resistances (31, 75).

One should note that many current interoceptive tasks may also be accomplished with ‘cognitive’ strategies. For example, heartbeat detection has been shown to be vulnerable to harmonization strategies where participants estimates of their heartbeats are based on their prior beliefs about their resting heart rate, as opposed to their momentary tracking of their own heart beats (76). It may be a difficult endeavor to expunge cognition from interoceptive tasks. Instead, we suggest future theory-driven use of control conditions. In the current paradigm, a control condition with paced counting divorced from breathing could isolate lapses related to working memory.

A final limitation of attentional paradigms writ large concerns the difficulty of temporally isolating attentional changes (e.g., lapses or meta-awareness of lapses). Some paradigms do not explicitly study prediction, instead examining periods of relatively decreased or increased attentional performance (19). Predictive paradigms typically involve forecasting responses to self-report probes during attentional tasks or resting state (20, 77). However, such probes—like the “cycle terminations” in our task—may reflect either the moment of attentional change or the moment of its detection, making temporal precedence difficult to establish. Emerging approaches, such as real-time triggered experience sampling in which physiological signals are monitored and used to trigger probes, may help better establish the temporal sequence of attentional changes (78).

## Conclusion

We conducted a novel examination of multimodal predictors of attentional lapses in breathing attention. Findings indicate generalizable between-individual predictors across response variability, breath variability, and attentional networks. At least in this task, there are indications that conventional top-down attentional systems are deployed to sustain breathing attention, along with interoceptive and somatosensory pathways. When supplemented by within-individual learning, the current multimodal markers could be used to facilitate attention during mind-body practices.

## Declarations

### Funding

This research was in part supported by NCCIH R01AT011002, NIMH R01MH116969, NIMH K23MH108752, the Tommy Fuss Fund, Klingenstein Third Generation Foundation, and a Young Investigator Grant from the Brain & Behavior Research Foundation awarded to CAW. Additionally, RPA received funding through the Tommy Fuss Fund, Dana Foundation, and Klingenstein Third Generation Foundation.

### Conflicts of interest/Competing interests

CAW has received consulting fees from King & Spalding law firm for unrelated work. In the past 3 years, RPA has received consulting fees and equity from Get Sonar Inc. He also has received consulting fees from RPA Health Consulting, Inc. and Covington & Burling LLP, which is representing a social media company in litigation. He also serves on the scientific advisory board for the Jake Collective. He also has received funding from AFSP, the Morgan Stanley Foundation, NIMH, and the Erick Shirley Foundation. All other authors report no biomedical financial interests or potential conflicts of interest.

### Ethics Approval

All procedures were approved by the Mass General Brigham IRB.

### Consent to Participate / Consent for Publication

We obtained informed consent / assent from all participants, including consent to having (deidentified) data published.

### Availability of data and materials

Data (predictors) will be available upon publication at https://osf.io/285xa/?view_only=db9e3408ac7e43ec9e039e313c9adb6b.

### Code availability

Code will be available upon publication at https://osf.io/285xa/?view_only=db9e3408ac7e43ec9e039e313c9adb6b.

### Author Contributions

INT contributed conceptualization, formal analysis, investigation, methodology, project administration, software, visualization, writing-original draft and writing-editing. CS contributed visualization, writing-editing, and software. AD contributed conceptualization, writing-original draft, and writing-editing. NJ and AOT contributed project administration and data curation. RPA contributed writing-editing. CAW contributed funding acquisition, methodology, project administration, writing-editing, resources, and supervision.

## Acknowledgements

We thank Aaron Kucyi for suggestions and support.

## Open Practices Statement

This study was preregistered at https://osf.io/fetvb. Data (predictors) and code will be available upon publication at https://osf.io/285xa/?view_only=db9e3408ac7e43ec9e039e313c9adb6b.

## Notes

### Summary of Updates

Introduction and Discussion are revised to better contextualize exteroceptive and interoceptive distinctions, highlight breath variability findings, and recognize limitations. A sensitivity analysis to GAD subjects is now included, as well as statistical tests of incremental validity.

