## Supplementary Information for "When attention falters: brain, breathing, and behavioral signals of lapses in interoceptive attention"

**Supplemental Materials** for **When attention falters: brain, breathing, and behavioral signals of lapses in interoceptive attention**

Isaac N. Treves^1,*^, Clare Shaffer^2^,Alexandra Decker^3^, Nigel Jaffe^4,5^, Anna O. Tierney^5^, Randy P. Auerbach^1^, Christian Webb^4,5,*^

**Supplement Text 1: Previous Neuroimaging Studies on Breath-Counting**

Yue et al., 2020: Examines differences in functional connectivity for breath-counting to rest as a function of mindfulness training. Mindfulness training (as compared to active control) makes connectivity between BC and rest more similar within the networks of the DMN, CEN and SN. No relationships to behavior or mindfulness trait change are found. (1)

Treves et al. 2025a: Examines differences in dynamic brain states for breath-counting vs rest. There is an additional between-individual comparison of performance on BCT vs dynamic brain state duration, as well as an exploratory within-task analysis of dynamic states, with no test of generalizability. (2)

*For between-person analysis of breath-counting behavior compared to external criteria, see Treves et al., 2025b.(3)*

**Supplement Text 2: Deviations from Preregistration**

We made the following deviations:

- We did not examine postdiction (responses to attentional lapses) in the current manuscript to narrow the scope and focus on models with clear practical implications for prediction.
- We used cluster corrected t-tests to examine timepoints instead of adding all the data to a logistic model, given the high correlations between timepoints.
- The actual sample size was larger than preregistered, as we decided to incorporate a generalizability sample (held-out test set).
- We chose to focus on random forest classification instead of the NBS-Predict (4). This was based on a) the lack of nonlinear ML methods in NBS-Predict and b) the requirement of ‘connected components’. Random forest models have often shown superior predictive performance in brain connectome applications(5).

And omitted the following exploratory analyses:

- We did not examine dynamic connectivity given concerns about windowing.

**Supplement Text 3: Brain Connectivity Results from Fixed Windows**

We trained models with fixed windows of the average window length (36.89 s) on binary classification of correct cycles, miscounts and resets. Random forest models using the Brainnetome atlas resulted in generalizable models of correct cycles (held-out AUC=0.653, non-parametric *p* < 0.001*)* and miscounts (held-out AUC=0.589, *p* < 0.001), but not resets (*p* > 0.05). Results were similar when regressing out head motion. Random forest models using the Schaefer atlas resulted in similar relationships (AUCs= 0.638, 0.59), respectively. Areas where higher connectivity predicted more likelihood of miscounts included the DAN, VAN, and cerebellum (**Figure S3)**. Areas where higher connectivity predicted more likelihood of correct responses included subcortex and FPN, and somatomotor networks (but only for brainnetome, **Figure S4**). These results largely aligned with the main models, excepting the inconsistency of within-somatomotor connectivity relationships (which were universally related to correct performance in the main models).

**Supplement Text 4: Moderation by Generalized Anxiety Disorder**

41.9% of our sample reported generalized anxiety disorder. Given that anxiety affects monitoring and interoceptive saliency, we analyzed whether diagnoses moderated terminations and model performance in predicting those terminations. We conducted DeLong’s test to statistically compare prediction AUC for GAD and no GAD. Simple t-tests of frequencies were used to compare rates of each termination type. Anxious individuals showed better performance than non-anxious individuals, primarily due to fewer miscounts. Across all modalities, performance of predicting resets was higher. No FDR-correction was performed, but a conservative threshold of p < 0.01 was used to assign asterisks.

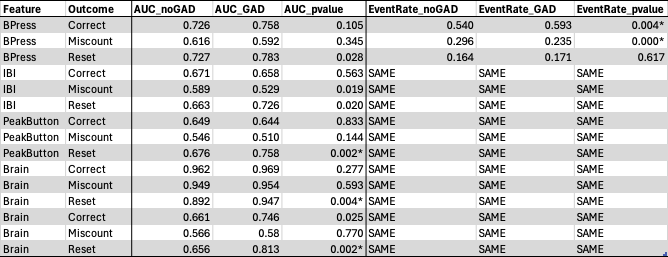

| *Networks* | *Model* | *Training Accuracy (Balanced)* | *Training set permutation p* | *Test set significant?* |
| --- | --- | --- | --- | --- |
| Brainnetome | Correct or Not | 0.552 | 0 | NULL |
| Brainnetome | Miscount or Not | 0.572 | 0 | NULL |
| Brainnetome | Reset or Not | 0.553 | 0 | NULL |
| Schaefer | Correct or Not | 0.52 | 0.31 | NULL |
| Schaefer | Miscount or Not | 0.573 | 0 | NULL |
| Schaefer | Reset or Not | 0.549 | 0 | NULL |

**Supplement Table 1:** NBS Toolbox results, summary. Networks: parcellations used for deriving connectivity features. Model: binarized outcomes. Training accuracy: model accuracy in training set. Permutation p: p-value based on shuffling labels 1000 times, values of zero reflect no permutations with higher accuracies. Test set significant: whether the model generalized to held-out participants. NULL: failure to reject null hypothesis.

| **Sample Characteristics** | | | |
| --- | --- | --- | --- |
|  | **N (*n =* 93)** |  | **%** |
| **Biological Sex** |  |  |  |
| Female | 73 |  | 78.5 |
| Male | 20 |  | 21.5 |
| **Race** |  |  |  |
| AI/AN | 1 |  | 1.1 |
| Asian | 14 |  | 15.1 |
| Black or African American | 4 |  | 4.3 |
| NH/OPI | 0 |  | 0.0 |
| White | 59 |  | 63.4 |
| Multiracial | 12 |  | 12.9 |
| Other | 3 |  | 3.2 |
| **Ethnicity** |  |  |  |
| Hispanic or Latino | 10 |  | 10.8 |
| Not Hispanic or Latino | 83 |  | 89.2 |
| **Current Diagnoses (DSM-V)** |  |  |  |
| MDD | 8 |  | 8.6 |
| GAD | 39 |  | 41.9 |
| SAD | 18 |  | 19.4 |
| Panic Disorder | 7 |  | 7.5 |
| Specific Phobia | 6 |  | 6.5 |
| Selective Mutism | 1 |  | 1.1 |
| ODD | 3 |  | 3.2 |
| OCD | 2 |  | 2.2 |
| Binge Eating Disorder | 3 |  | 3.2 |
| PTSD | 1 |  | 1.1 |
| Alcohol / Substance Use Disorders | 0 |  | 0.0 |
| **Handedness** |  |  |  |
| Right | 63 |  | 67.7 |
| Left | 5 |  | 5.4 |
| Unknown | 25 |  | 26.9 |
|  | **M (Range)** |  | **SD** |
| **Age (years)** | 15.7 (13 -18) |  | 1.7 |
| **Family Income (dollars)** | 168,293.8 (0 - 600,000) |  | 132,937.9 |

**Supplement Table 2.** Demographic and clinical characteristics of the sample. AI/AN: American Indian or Alaska Native. NH/OPI: Native Hawaiian or Other Pacific Islander. MDD: Major Depressive Disorder. GAD: Generalized Anxiety Disorder. SAD: Social Anxiety Disorder. ODD: Oppositional Defiant Disorder. OCD: Obsessive Compulsive Disorder. PTSD: Post-Traumatic Stress Disorder. SUDs = Substance Use Disorders.

| Predictor | Outcome | Feature | Effect (Beta) | P_value | 95% CI |
| --- | --- | --- | --- | --- | --- |
| ButtonPress | Correct | CV | -1.019* | 0.0000 | [-1.183, -0.854] |
| ButtonPress | Correct | MeanLevel | -0.263* | 0.0032 | [-0.437, -0.088] |
| ButtonPress | Miscount | CV | 0.468* | 0.0000 | [0.341, 0.596] |
| ButtonPress | Reset | CV | 0.484* | 0.0000 | [0.328, 0.639] |
| ButtonPress | Reset | MeanLevel | 0.543* | 0.0000 | [0.316, 0.77] |
| IBI | Correct | CV | -0.183* | 0.0035 | [-0.305, -0.06] |
| PeakButton | Correct | CV | -0.243* | 0.0004 | [-0.376, -0.109] |
| PeakButton | Correct | Slope_Predictor | 0.211* | 0.0017 | [0.079, 0.342] |
| PeakButton | Miscount | CV | 0.203* | 0.0043 | [0.064, 0.342] |
| PeakButton | Miscount | Slope_Predictor | 0.221* | 0.0031 | [0.074, 0.368] |
| PeakButton | Reset | MeanLevel | -0.326* | 0.0087 | [-0.569, -0.082] |
| PeakButton | Reset | Slope_Predictor | -0.588* | 0.0000 | [-0.766, -0.41] |

**Supplement Table 3:** Significant features in training set logistic models for longer window. Only features meeting strict Bonferroni threshold are displayed. Features were computed over the long time window (50 seconds), excluding any overlap with previous cycles. Betas are standardized. CV: coefficient of variation of predictor, MeanLevel: overall mean of predictor in time window, Slope_Predictor: linear slope of predictor.

| Predictor | Outcome | Feature | Effect (Beta) | P_value | 95% CI |
| --- | --- | --- | --- | --- | --- |
| ButtonPress | Correct | CV | -1.003* | 0.0000 | [-1.17, -0.837] |
| ButtonPress | Correct | MeanLevel | -0.391* | 0.0000 | [-0.564, -0.219] |
| ButtonPress | Miscount | CV | 0.398* | 0.0000 | [0.274, 0.522] |
| ButtonPress | Reset | CV | 0.481* | 0.0000 | [0.327, 0.635] |
| ButtonPress | Reset | MeanLevel | 0.681* | 0.0000 | [0.466, 0.896] |
| IBI | Reset | CV | 0.241* | 0.0019 | [0.089, 0.392] |
| PeakButton | Correct | CV | -0.223* | 0.0010 | [-0.356, -0.09] |
| PeakButton | Correct | Slope_Predictor | 0.195* | 0.0015 | [0.075, 0.316] |
| PeakButton | Reset | Slope_Predictor | -0.501* | 0.0000 | [-0.666, -0.337] |

**Supplement Table 4:** Significant features in training set logistic models for shorter time window. Only features meeting strict Bonferroni threshold are displayed. Features were computed over the short time window (25 seconds), excluding any overlap with previous cycles. Betas are standardized. CV: coefficient of variation of predictor, MeanLevel: overall mean of predictor in time window, Slope_Predictor: linear slope of predictor.

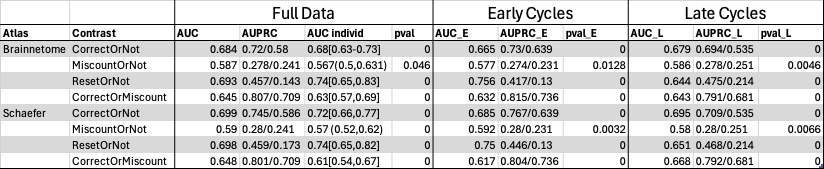

**Supplement Table 5:** Performance for Main Contrasts of Interest, Brain Connectivity Models. Brainnetome and Schaefer atlases show similar performance. Early and late cycles are first and second half of participant cycles, respectively. AUC: area under curve; AUPRC: area under precision-recall curve, sensitive to baseline frequency of positive class (shown as ratio), AUC individ: AUC distribution with 95% CI for each participant, *pval*: motion-adjusted AUC significance when shuffling outcome labels, where a *p*-val of 0 means none of the 1000 shuffles had higher AUCs, AUC_E: early AUC, AUC_L: late AUC. Individaul AUCs are not shown for early and late trials to simplify display. A noticeable fatigue effect may be noted for Resets, where early performance is > .1 AUC above late performance.

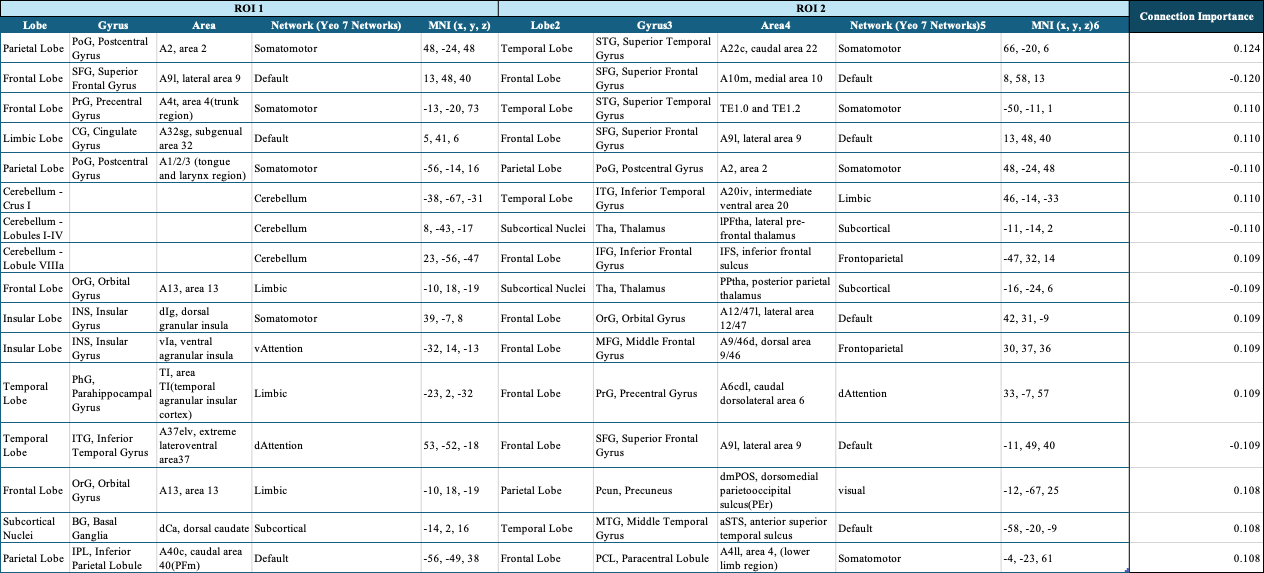

**Supplement Table S6.** Top 5% of Brainnetome Connections (RO1-ROI2) for models predicting correct vs miscount trials. Positive importances reflect higher likelihood of correct responses, whereas negative importances reflect higher likelihood of miscounts. Left columns show ROI1 information, and right columns show ROI2 information.

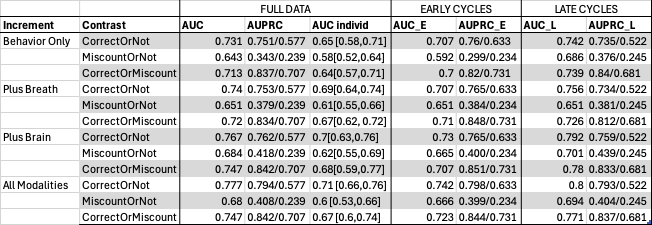

**Supplement Table S7.** Performance for Multimodal Models. Brainnetome and Schaefer atlases show similar performance- here Schaefer is shown. Behavior only refers to just button pressing features; Plus Breath adds inter-breath-interval features; Plus Brain adds connectivity; and All Modalities is all three. All models were significant. AUCs appear to be slightly higher when incorporating brain data. Early and late cycles are first and second half of participant cycles, respectively. AUC: area under curve; AUPRC: area under precision-recall curve, sensitive to baseline frequency of positive class (shown as ratio), AUC individ: AUC distribution with 95% CI for each participant, AUC_E: early AUC, AUC_L: late AUC. Individual AUCs are not shown for early and late trials to simplify display.

SUBJECT 116

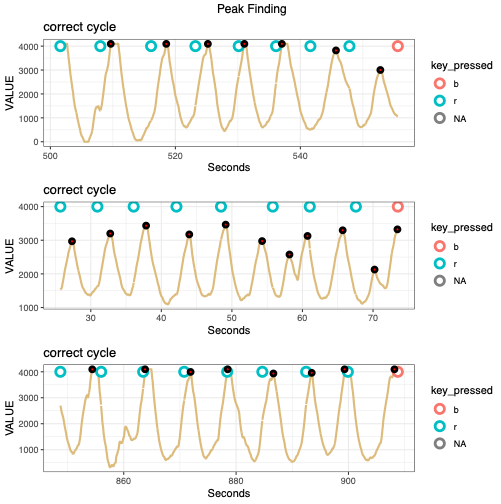

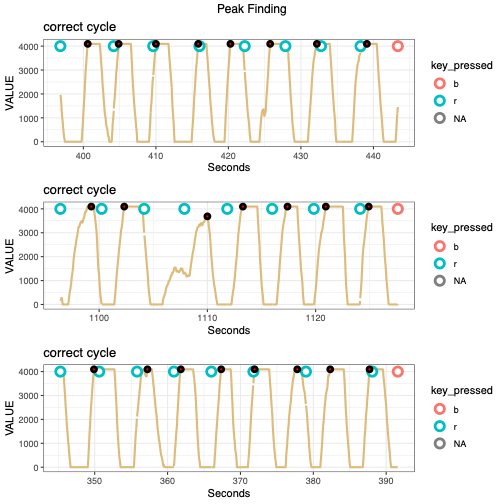

SUBJECT 102

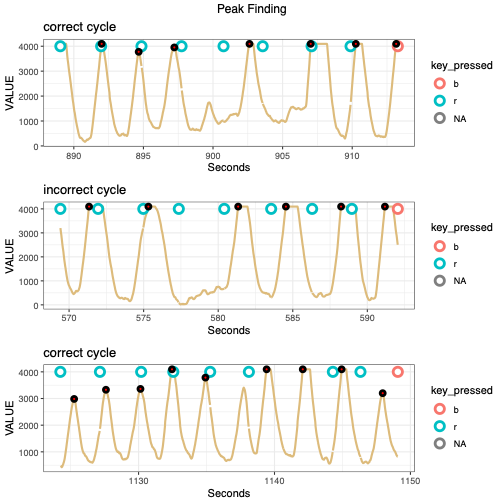

SUBJECT 111

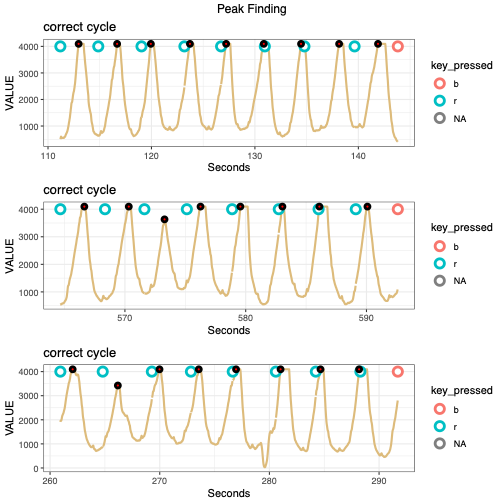

SUBJECT 183

**Figure S1:** Peak finding QC for four subjects for randomly chosen cycles. Respiratory traces are shown in orange. Detected peaks are indicated with small black circles. The algorithm used is : *pracma::findpeaks(DataPhysio$VALUE,minpeakheight=2000, minpeakdistance=100,nups=2,ndowns=2,zero="+"),* with the exception of 134 and 157 who used custom algorithms.

SUBJECT 183

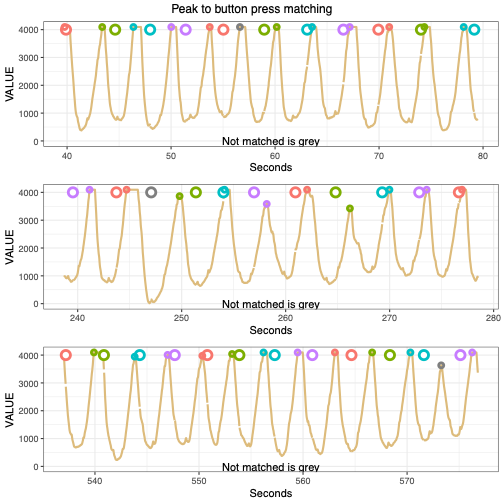

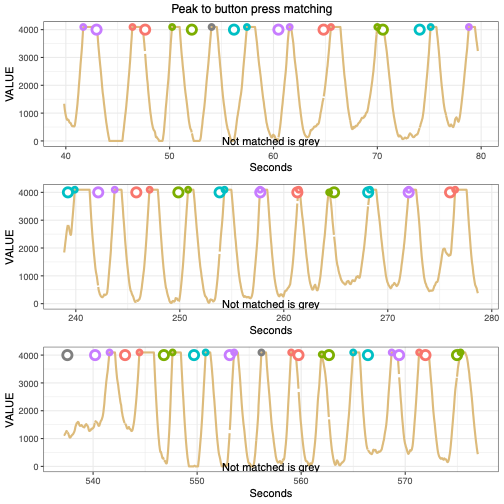

SUBJECT 111

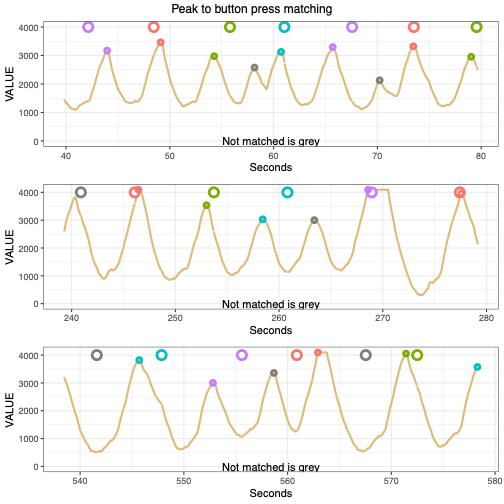

SUBJECT 116

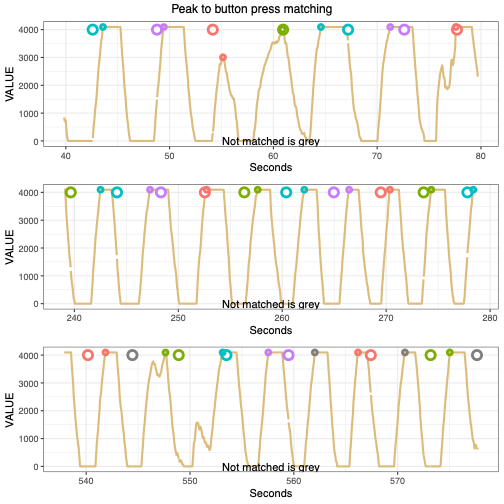

SUBJECT 102

**Figure S2:** Peak button press matching QC for four subjects for randomly chosen cycles. The closest function in R was used, which finds unique matches that minimize the distances between two discrete vectors. Respiratory traces are shown in orange. Detected peaks are indicated with small circles, with colors matching the corresponding button press. Detected peaks or button presses that were not matched are in grey.

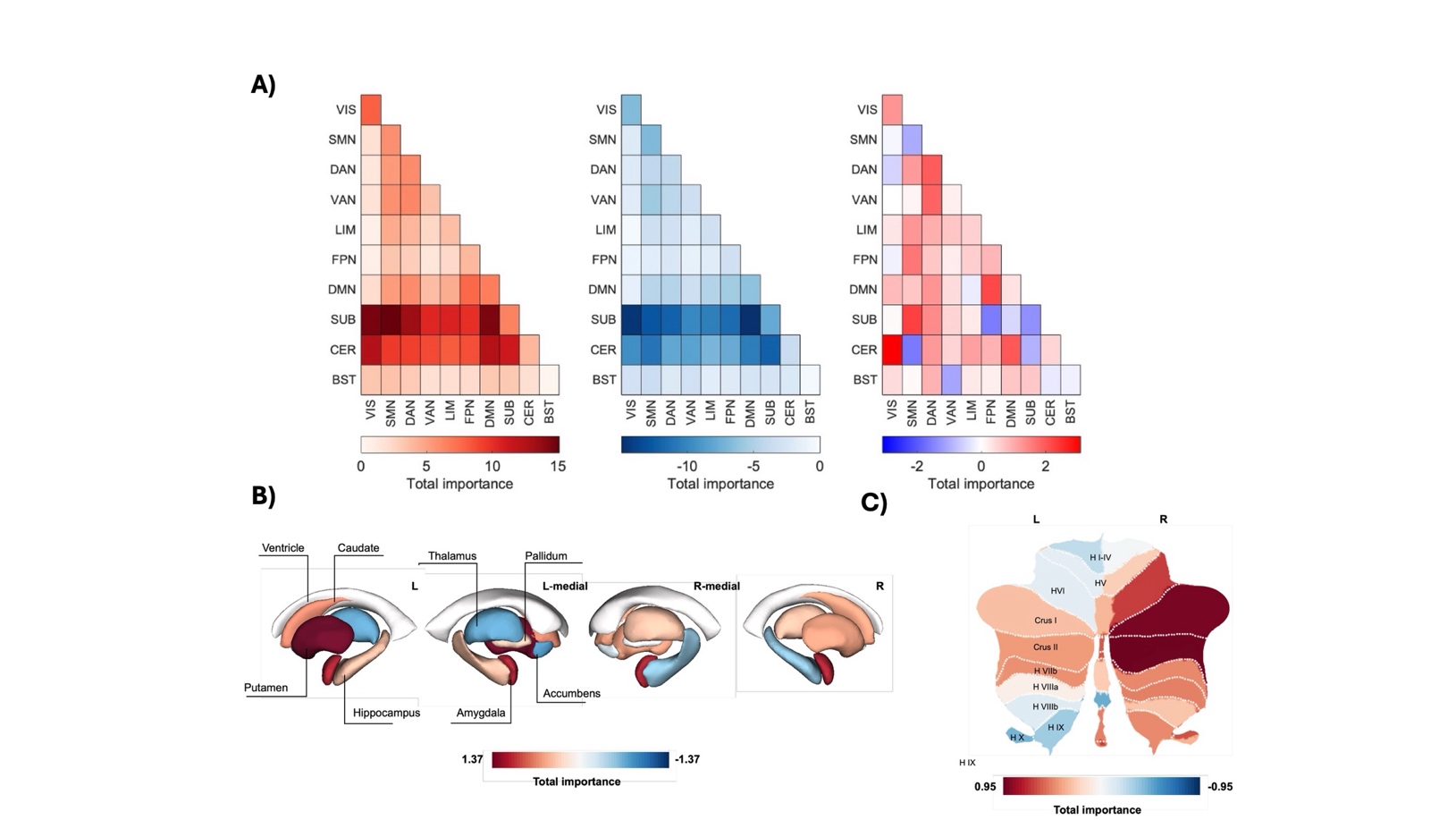

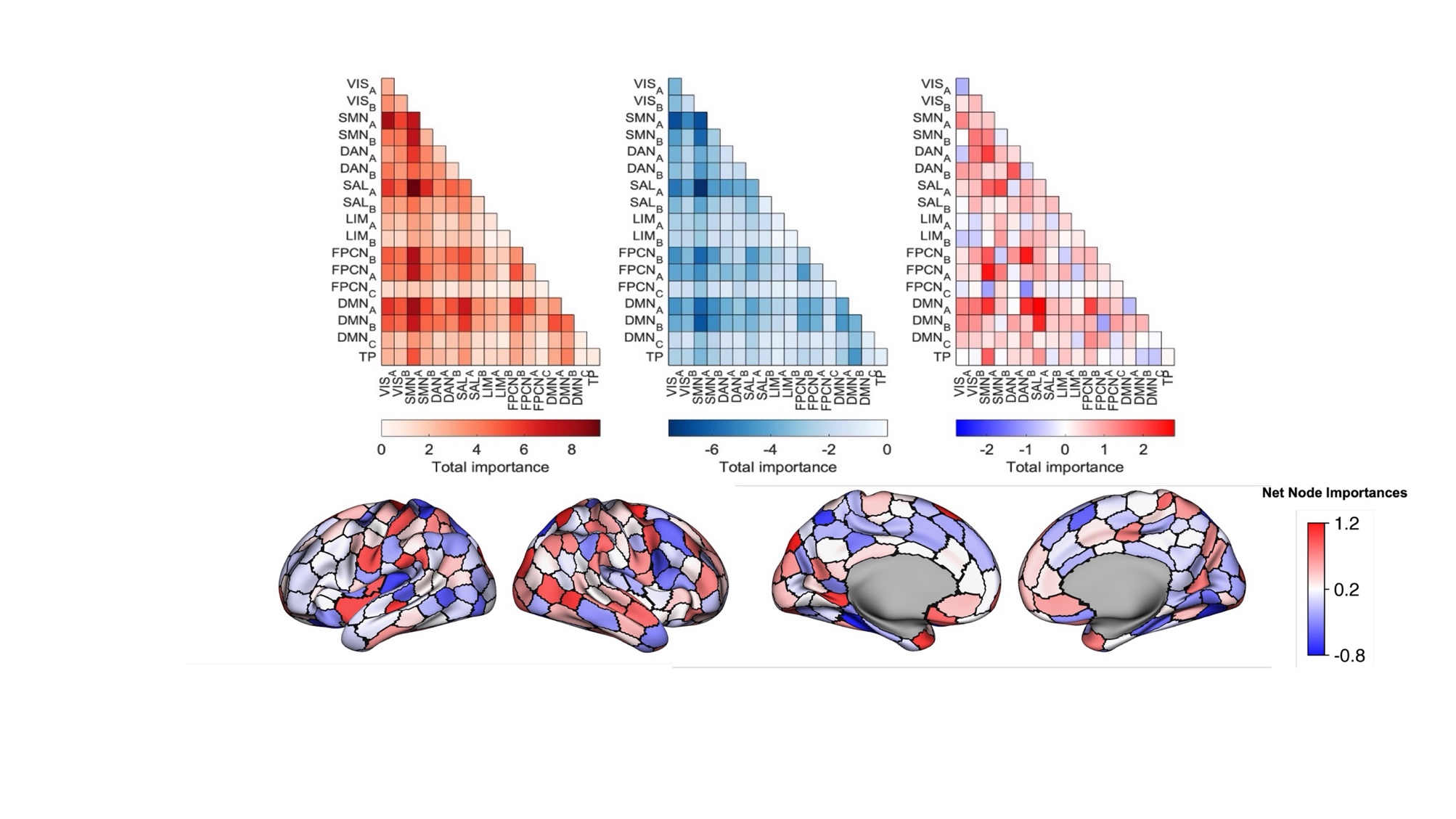
**Figure S3:** Fixed window (36.89 s) connectivity models to predict miscounts, Brainnetome whole-brain atlas. More positive edges predict miscounts, whereas negative edges predict correct responses. The left-most panel is restricted to positive edges only, the middle panel is restricted to negative edges only, and the right-most panel is the net importance.

**Figure S4:** Fixed window (36.89 s) connectivity models to predict miscounts, Schaefer cortical atlas. More positive edges predict miscounts, whereas negative edges predict correct responses. The left-most panel is restricted to positive edges only, the middle panel is restricted to negative edges only, and the right-most panel is the net importance

**Figure S5:** Held-out performance of button press models for attentional lapses. On left, permutation significance distributions (shuffling labels), with the true AUC showed in red. All p-values are less than <0.001. On right, ROC Curves with AUCs.

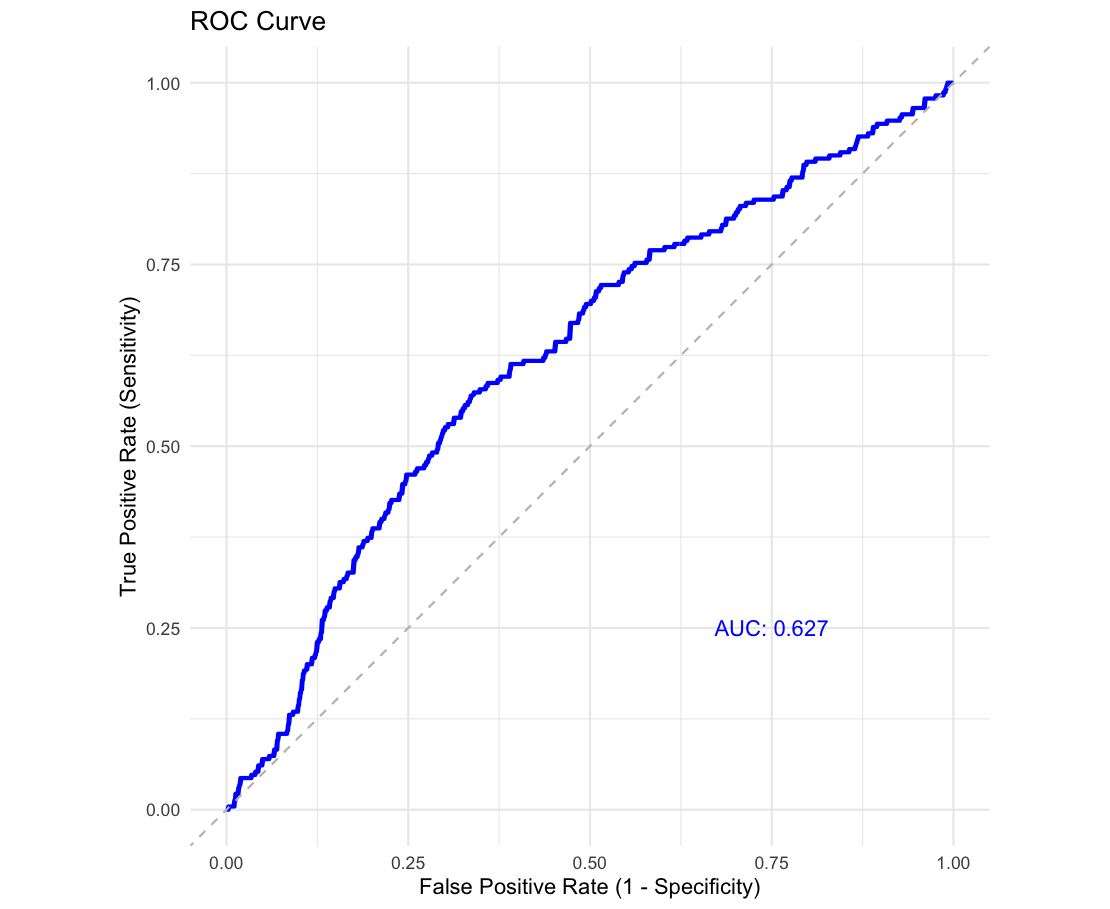

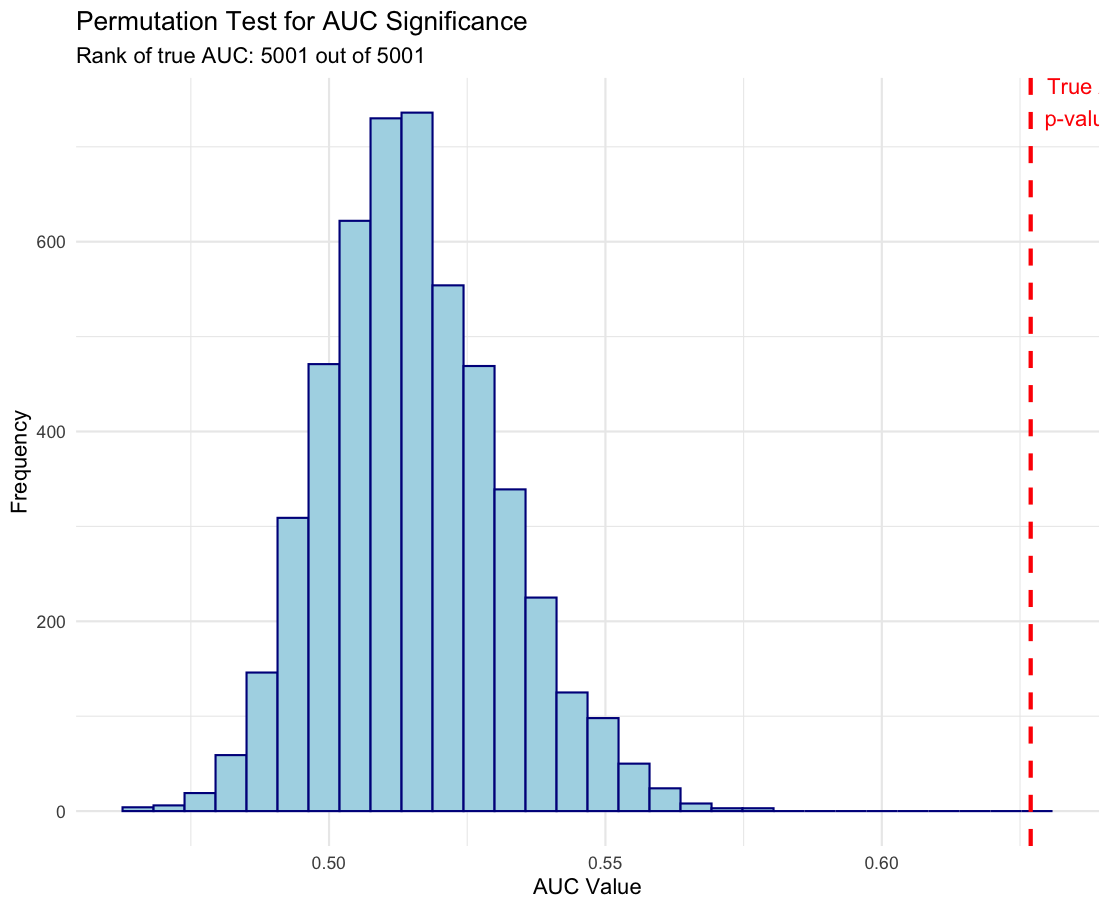

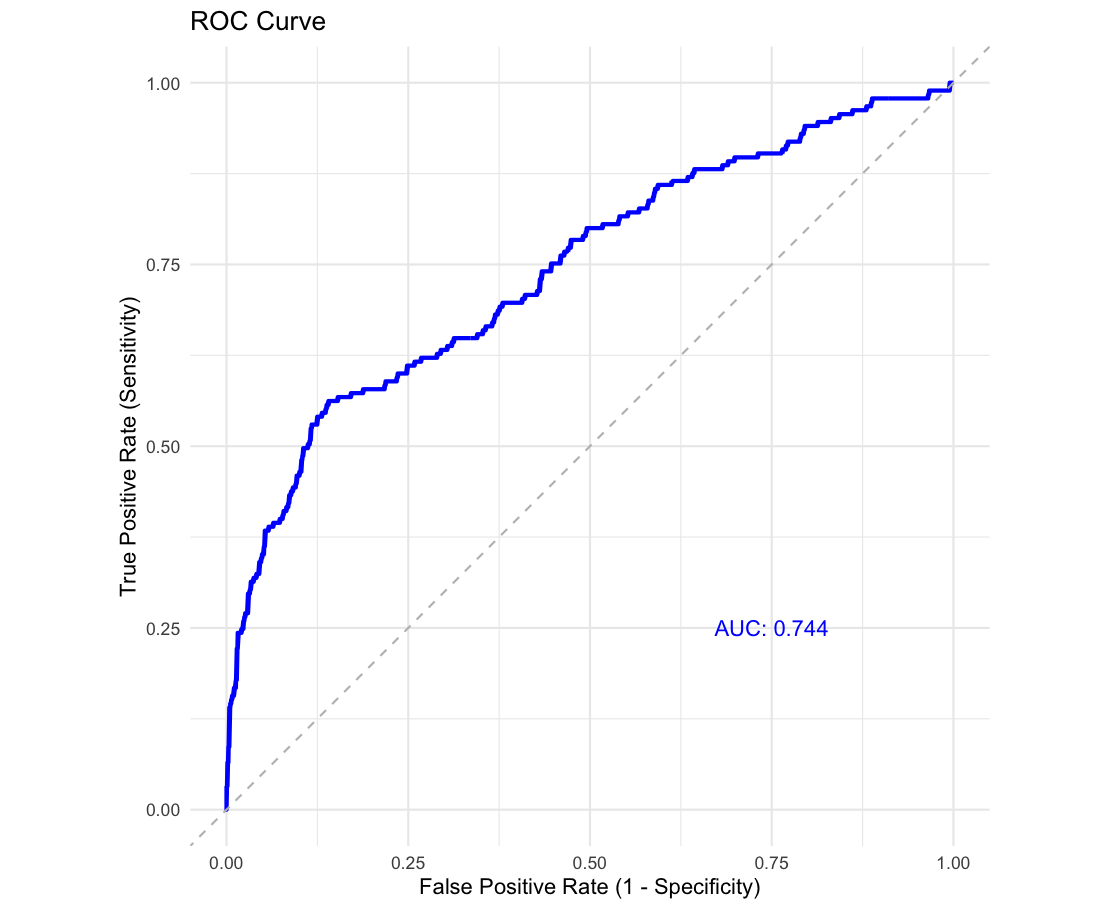

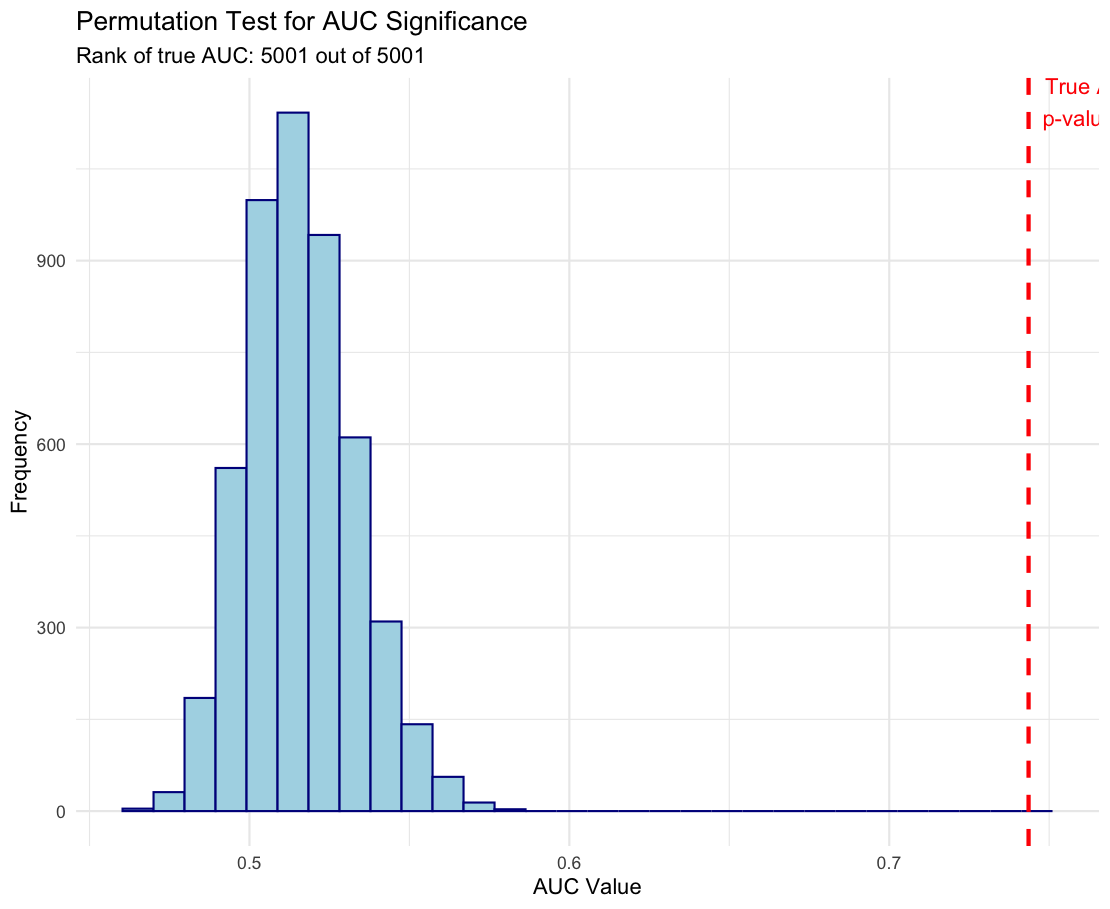

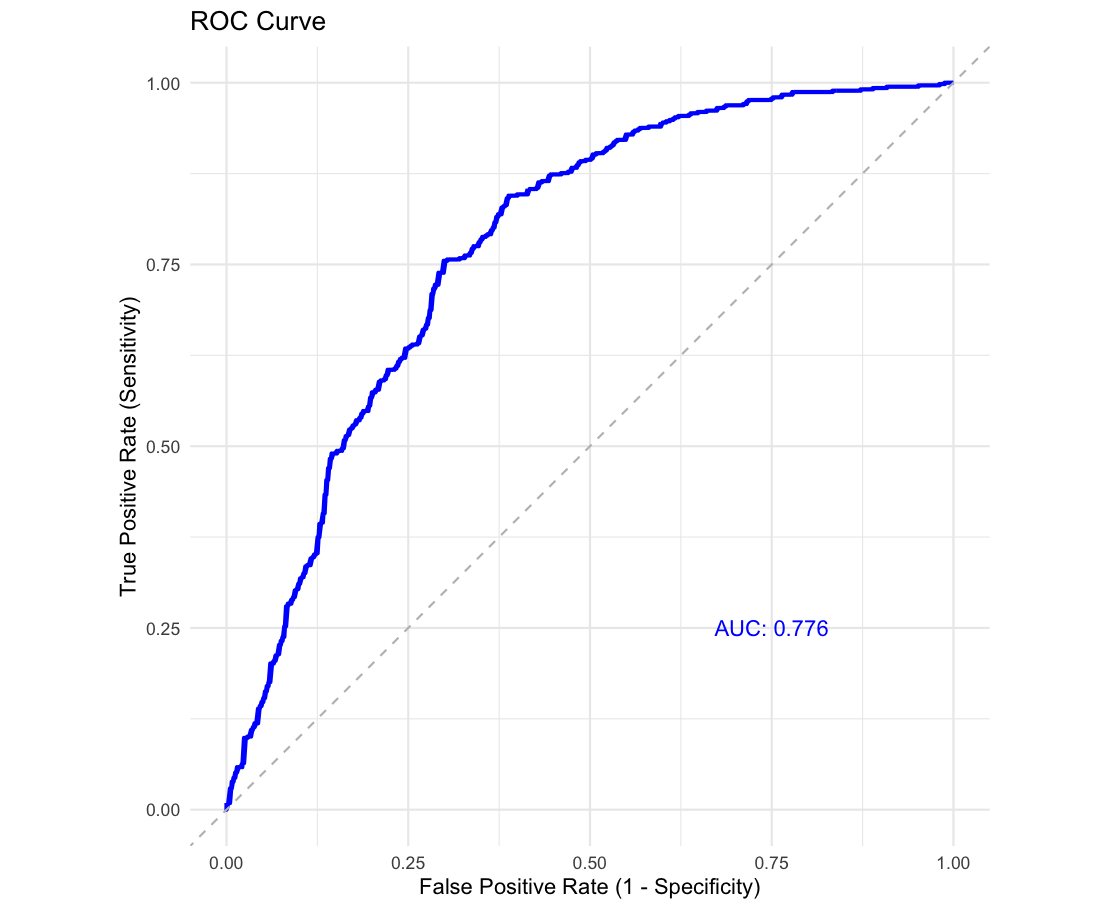

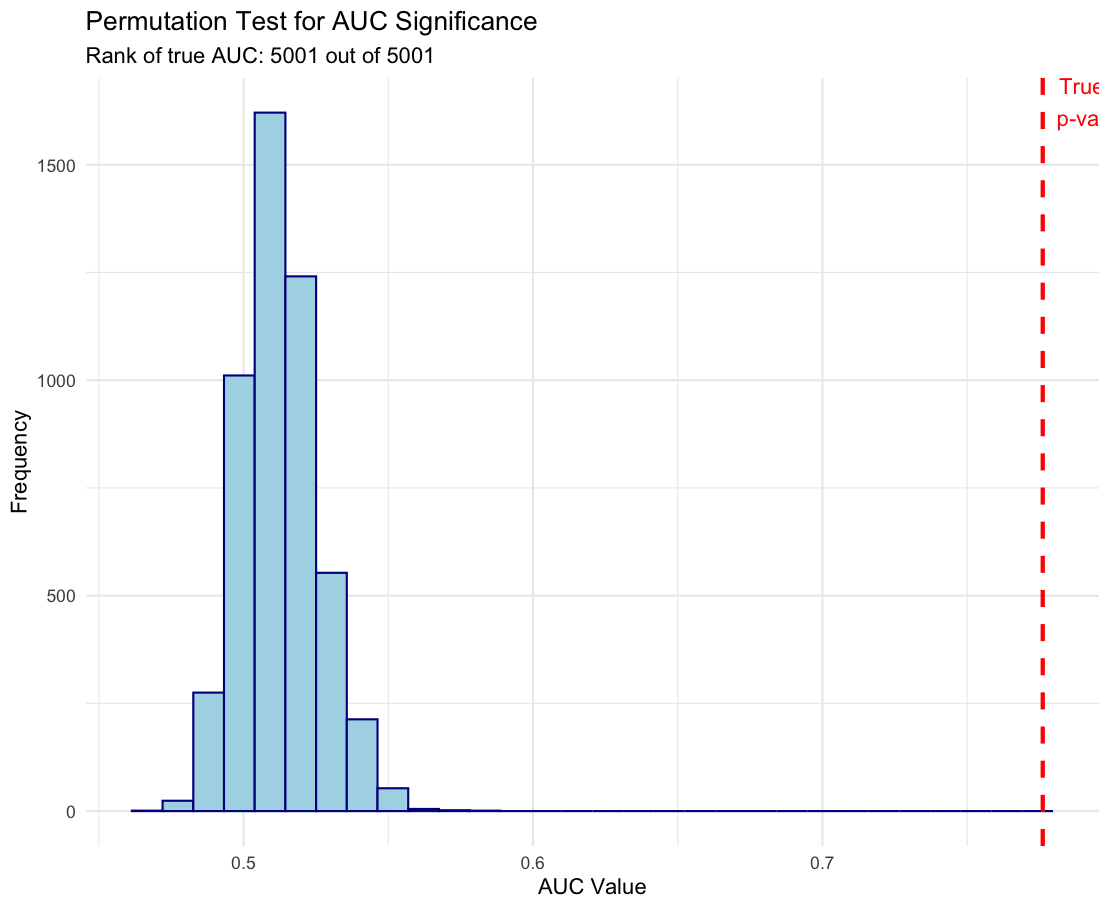

**A.Correct**

**B. Reset**

**C. Miscount**

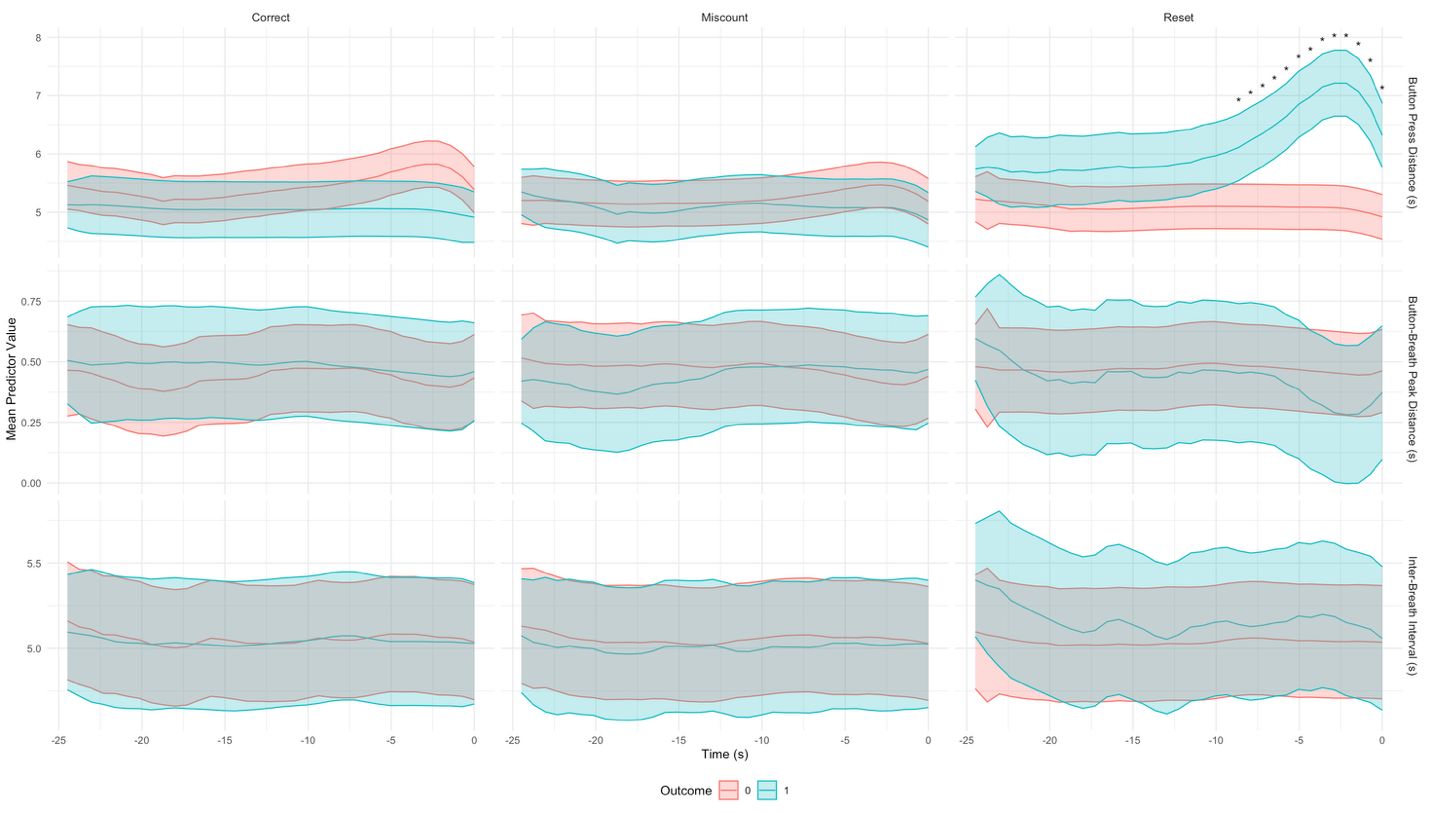

**Supplement Figure 6:** Mean predictive values of behavioral and physiological predictors are shown across individuals for shorter window size of 25 seconds. Timepoints (x-axis) are shown preceding the termination of a cycle (at 0). Predictive values are shown separately for correct responses (left column), miscounts (middle column), and resets (right columns). Each row reflects a distinct type of predictor (BPI, top; BBI (middle) IBI (bottom). Blue represents the presence of the termination (e.g., ‘correct’), red is an alternative termination (e.g., not ‘correct’) . Lines reflect means from multilevel models with categorical time variable to allow for nonlinearities, shading reflects 95% CI. Cycle number is included in models to account for fixed effect of fatigue. Asterisks indicate non-overlapping confidence intervals.

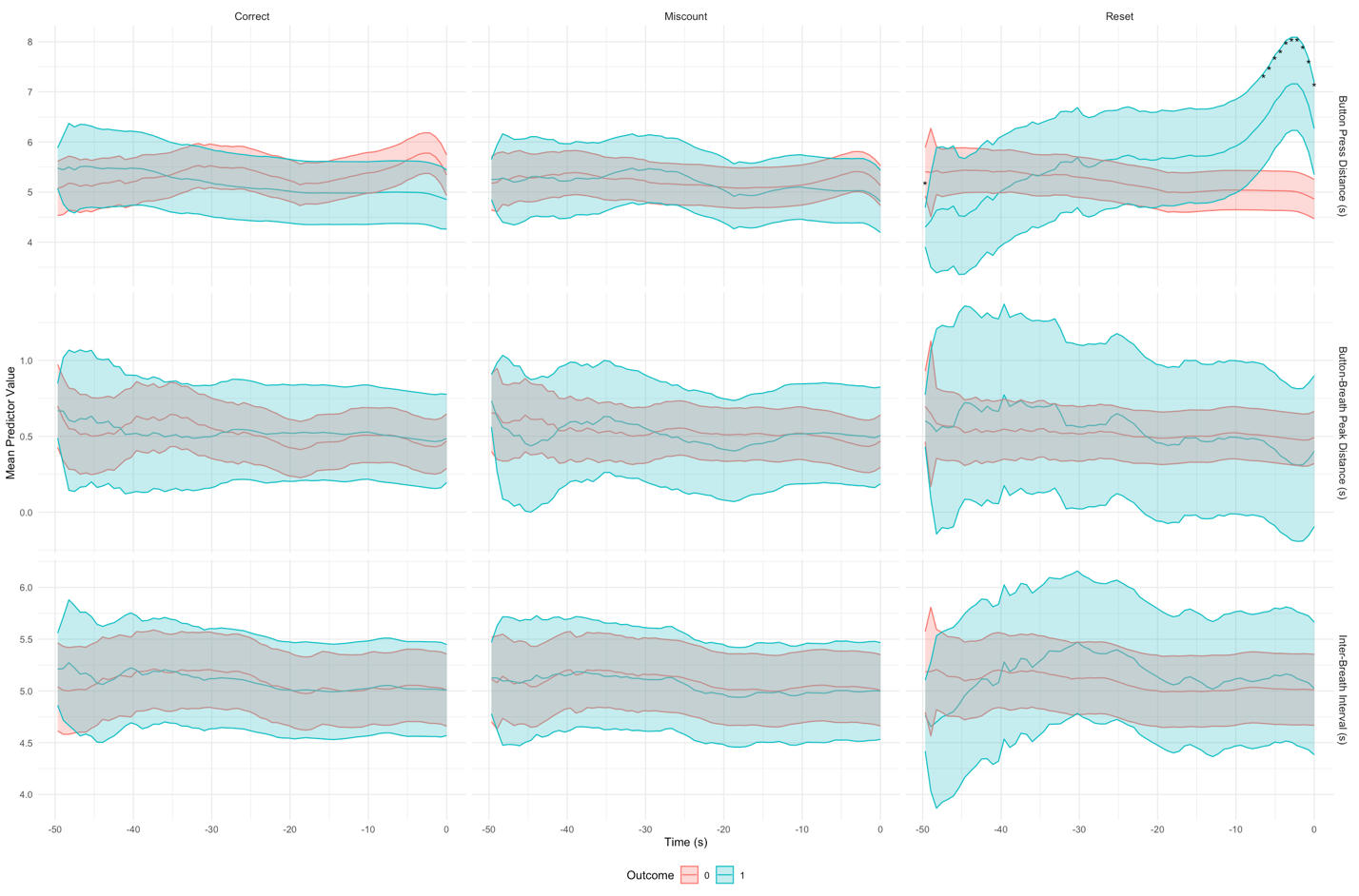

**Supplement Figure 7:** Mean predictive values of behavioral and physiological predictors are shown across individuals for longer window size (50s). Timepoints (x-axis) are shown preceding the termination of a cycle (at 0). Predictive values are shown separately for correct responses (left column), miscounts (middle column), and resets (right columns). Each row reflects a distinct type of predictor (BPI, top; BBI (middle) IBI (bottom). Blue represents the presence of the termination (e.g., ‘correct’), red is an alternative termination (e.g., not ‘correct’) . Lines reflect means from multilevel models with categorical time variable to allow for nonlinearities, shading reflects 95% CI. Cycle number is included in models to account for fixed effect of fatigue. Asterisks indicate non-overlapping confidence intervals.

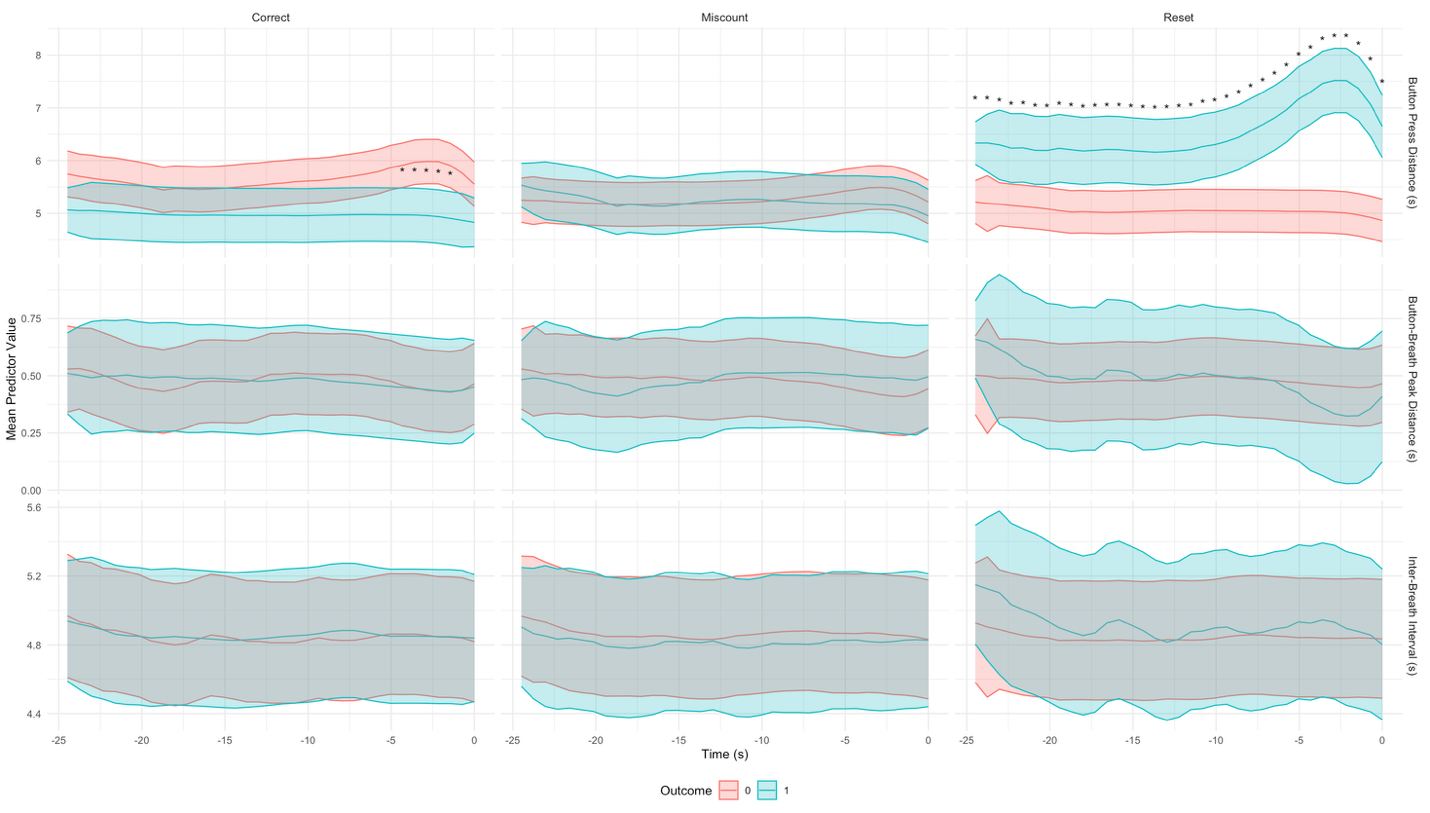

**Supplement Figure 8:** Mean predictive values of behavioral and physiological predictors are shown across individuals, without controlling for fatigue. Timepoints (x-axis) are shown preceding the termination of a cycle (at 0). Predictive values are shown separately for correct responses (left column), miscounts (middle column), and resets (right columns). Each row reflects a distinct type of predictor (BPI, top; BBI (middle) IBI (bottom). Blue represents the presence of the termination (e.g., ‘correct’), red is an alternative termination (e.g., not ‘correct’) . Lines reflect means from multilevel models with categorical time variable to allow for nonlinearities, shading reflects 95% CI. Asterisks indicate non-overlapping confidence intervals.

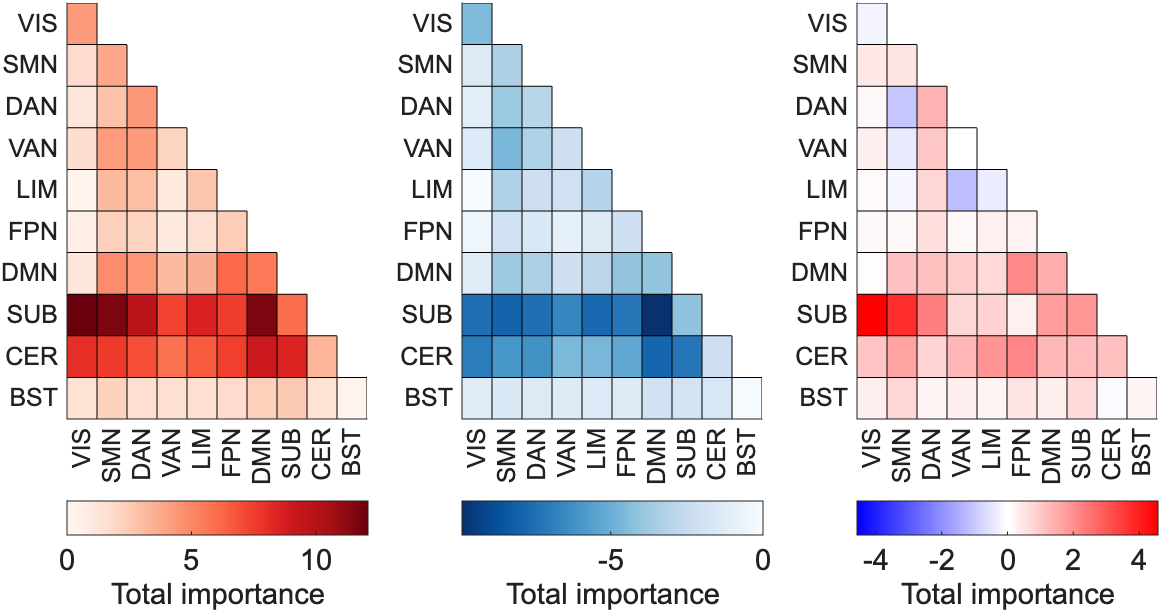

**Figure S9:** Connectivity models to predict resets vs non-resets, Brainnetome atlas. More positive edges predict cycles where participants lost count and reset the cycle, whereas negative edges predict all other cycle types. The left-most panel is restricted to positive edges only, the middle panel is restricted to negative edges only, and the right-most panel is the net importances. Net importances show particularly strong positive associations with resets are found in subcortical-VIS and subcortical-SMN connectivity.

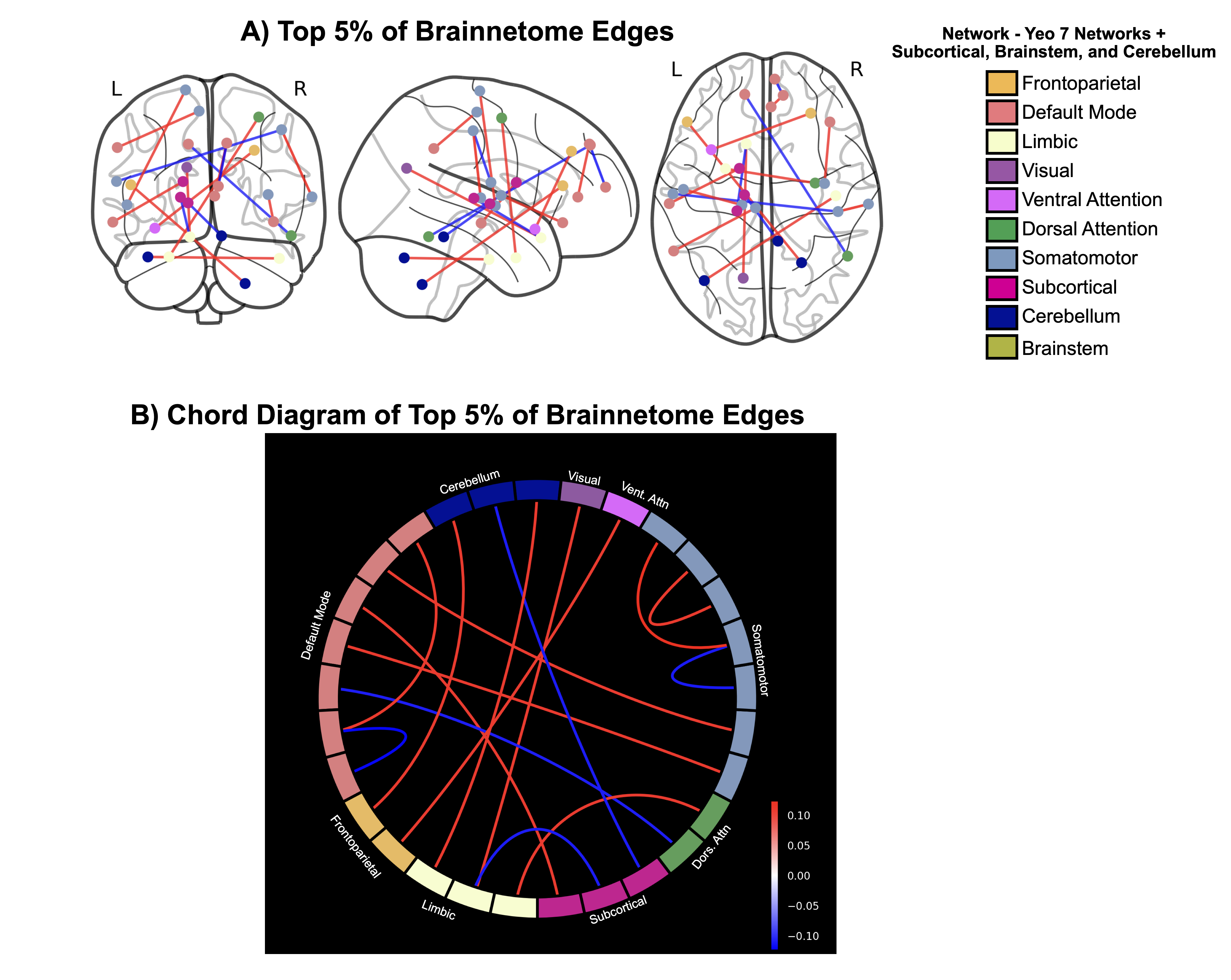

**Supplement Figure 10.** Top 5% of connections in Brainnetome models predicting miscounts versus correct trials. Positive importances reflect higher likelihood of correct responses, whereas negative importances reflect higher likelihood of miscounts. A) Connections are plotted on a glass brain to depict the contributions of individual nodes and their connections. Blue lines represent negative weights; red lines represent positive weights. B) Importance weights for each of the top 5% of connections are plotted on a chord diagram, where different contributions from the same network are shown. For miscount trials, multiple networks contribute multiple connections to the top 5% of importance weights.

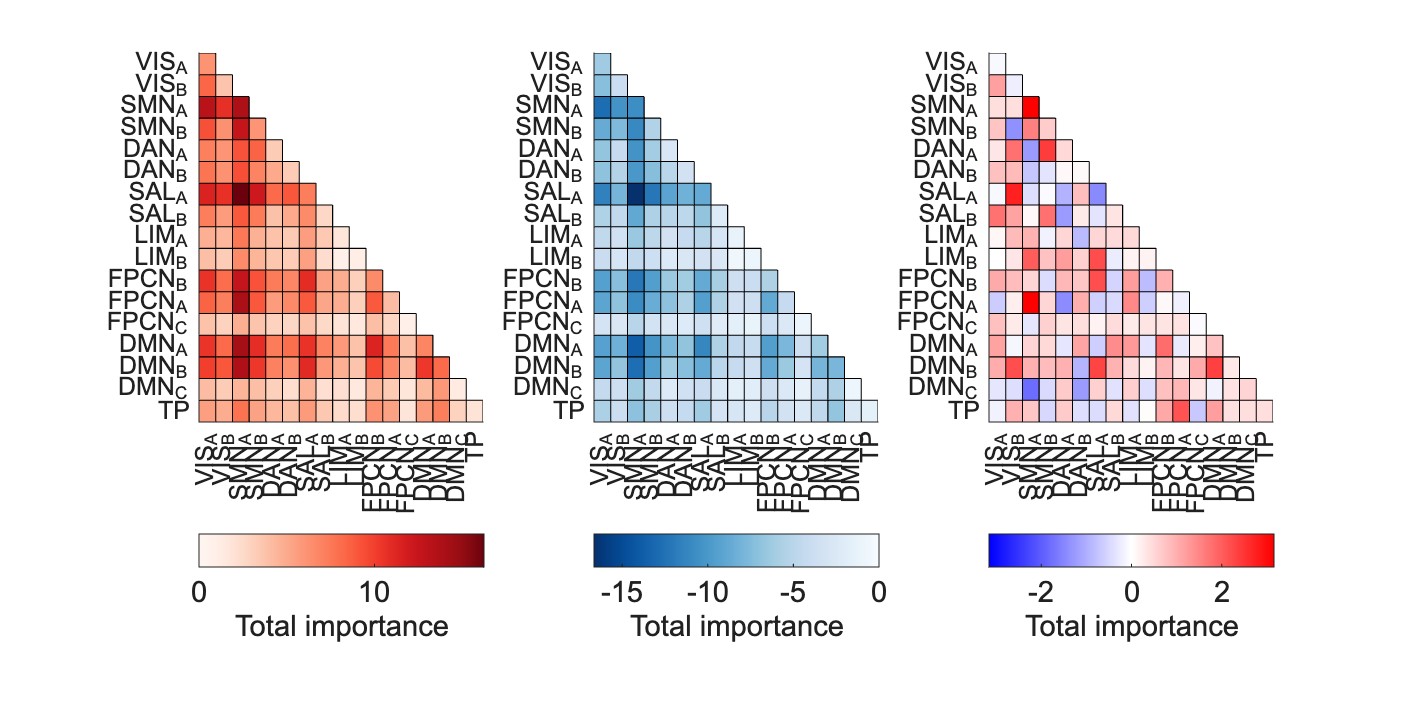

**Figure S11:** Connectivity models to predict correct vs incorrect breathing, Schaefer cortical atlas. More positive edges predict cycles where breathing corresponds to counts (correct breathing), whereas negative edges predict discordant cycles (incorrect breathing). The left-most panel is restricted to positive edges only, the middle panel is restricted to negative edges only, and the right-most panel is the net importance.
